# Evaluation of high-resolution atmospheric and oceanic simulations of the California Current System

**DOI:** 10.1101/2020.02.10.942730

**Authors:** Lionel Renault, James C. McWilliams, Faycal Kessouri, Alexandre Jousse, Hartmut Frenzel, Ru Chen, Curtis Deutsch

## Abstract

This paper is the first of two that present a 16-year hindcast solution from a coupled physical and biogeochemical model of the California Current System (CCS) along the U. S. West Coast and validate the physical solution with respect to mean, seasonal, interannual, and sub-seasonal fields and, to a lesser degree, eddy variability. Its companion paper is Deutsch et al. (2021a). The intent is to construct and demonstrate a modeling tool that will be used for mechanistic explanations, attributive causal assessments, and forecasts of future evolution for circulation and biogeochemistry, with particular attention to the increasing oceanic stratification, deoxygenation, and acidification. A well-resolved mesoscale (*dx* = 4 km) simulation of the CCS circulation is made with the Regional Oceanic Modeling System over a hindcast period of 16 years from 1995 to 2010. The oceanic solution is forced by a high-resolution (*dx* = 6 km) regional configuration of the Weather and Research Forecast (WRF) atmospheric model. Both of these high-resolution regional oceanic and atmospheric simulations are forced by lateral open boundary conditions taken from larger-domain, coarser-resolution parent simulations that themselves have boundary conditions from the Mercator and Climate Forecast System reanalyses, respectively. We show good agreement between the simulated atmospheric forcing of the oceanic and satellite measurements for the spatial patterns and temporal variability for the surface fluxes of momentum, heat, and freshwater. The simulated oceanic physical fields are then evaluated with satellite and *in situ* measurements. The simulation reproduces the main structure of the climatological upwelling front and cross-shore isopycnal slopes, the mean current patterns (including the California Undercurrent), and the seasonal, interannual, and subseasonal variability. It also shows agreement between the mesoscale eddy activity and the windwork energy exchange between the ocean and atmosphere modulated by influences of surface current on surface stress. Finally, the impact of using a high frequency wind forcing is assessed for the importance of synoptic wind variability to realistically represent oceanic mesoscale activity and ageostrophic inertial currents.

## 1 Introduction

Subtropical eastern boundary upwelling systems like the California Current System (CCS) are among the biologically most productive coastal environments (Carr and Kearns, 2003), supporting some of the world’s major fisheries (FAO, 2009). Seasonal upwelling (mainly during spring and summer) of deep nutrient-rich water maintains high rates of productivity over broad scales (*e.g*., Chavez and Messie (2009)). Additionally, coastal currents and oceanic mesoscale variability contribute to cross-shore exchange of heat, salt, and biogeochemical materials between the open and coastal oceans as well as the surface layer and interior (Hickey, 1998; Capet et al., 2008a; Gruber et al., 2011; Renault et al., 2012, 2016a).

The seasonal upwelling introduces water with low dissolved oxygen and low pH (*i.e*., a below-critical carbonate saturation state) into the surface waters (Chan et al., 2008; Feely et al., 2008; Gruber et al., 2012), making this region more prone to hypoxia and acidification (Feely et al., 2018; Gruber et al., 2012). Shoaling of deep, low-oxygen and low-pH waters is particularly pertinent in the CCS because the eastern Pacific Ocean contains the world’s largest mid-depth Oxygen Minimum Zone (OMZ). This OMZ has been expanding (e.g., Stramma et al. (2010)), which makes the coast more susceptible to hypoxic intrusions onto its narrow continental shelf (*e.g*., Pennington et al. (2006)). The entire CCS is subject to large-scale climate changes (e.g., Stramma et al. (2012); Bopp et al. (2015); Garcia-Reyes et al. (2015)) that include deoxygenation caused in part by increased density stratification through anomalous greenhouse heating and the acidification due to anthropogenic CO_2_ invasion. These global influences may be exacerbated in coastal regions by pollutants deposited from the atmosphere (e.g., nitric and sulfuric acid), from urban wastewater effluents, and from riverine eutrophication and other contaminants. Multi-decadal declines in oxygen leading to hypoxia have also been observed in the coastal water off southern California and Oregon and have altered the proportions of biologically important nutrients. Dramatic responses to these perturbations have already been observed in species that form critical links in the food web (*e.g*., pteropods for oceanic acidification (Bednarsek and Ohman, 2015; Bednarsek et al., 2017) and benthic species and anchoviesfor hypoxia (Grantham et al., 2004; Feely et al., 2018; Howard et al., 2020b)).

Regional upwelling patterns and eddies are important influences on the ecosystem. Mesoscale eddies, induced by baroclinic and barotropic instabilities of the wind-driven currents (*e.g*., Marchesiello et al. (2003)), are present everywhere in the world ocean and play a key role in many oceanic processes. Many studies have shown their crucial role in the transport of heat and freshwater (e.g., Wunsch (1999); Dong et al. (2014)) and of biogeochemical materials (McGillicuddy, 2016). In the open ocean, mesoscale processes can enhance the biological production by increasing the surface concentration of limiting nutrients (McGillicuddy, 2016). In eastern boundary upwelling systems, eddies are a limiting factor that reduces the autotrophic primary production by fluxing unconsumed surface nutrients beneath the euphotic layer (“eddy quenching”) (Gruber et al., 2011; Renault et al., 2016a). As shown by Renault et al. (2016a,b); Desbiolles et al. (2016), a realistic representation of the slackening of the wind toward the coast (*i.e*., wind drop-off) is influential for the mean and mesoscale currents, the primary production, and interior oxygen levels (Deutsch et al., 2021b).

Equilibrium regional oceanic circulation models have been successfully employed for more than a decade in the CCS. As detailed briefly hereafter, many of the previous modeling efforts allowed a breakthrough in the understanding and modeling of the CCS. For instance, Marchesiello et al. (2003) was one of the first realistic mesoscale resolving regional simulation of the CCS; they forced a Regional Oceanic Modeling System (ROMS) simulation using climatological forcing derived from COADS. Veneziani et al. (2009) evaluate favorably with respect to measurements a high-resolution ROMS oceanic simulation forced by an interannual atmospheric forcing derived from COAMPS. Based on the same configuration, Neveu et al. (2016) successfully assimilate in *situ* and satellite data and fairly reproduce the CCS circulation and its main characteristics. Seo et al. (2016) and Renault et al. (2016d) couple a high-resolution oceanic simulation to a high resolution atmospheric simulation for a period of ≈ 5% years of the CCS. They show the large impact of air-sea interactions on the mesoscale activity. More recently, Fiechter et al. (2018) uses a *dx* = 3 km solution coupled with a biogeochemical model forced by the CCMP winds to assess the modulation of phytoplankton variability by the wind, the oceanic circulation, and topographic effects. From these studies, some of the oceanic simulations assimilate data, some other are coupled with the atmosphere or use interannual atmospheric forcing. However, no previous simulation has been made over a long time period using high-resolution spatial and temporal atmospheric forcing that includes the effects of wind drop-off (*i.e*., the cross-shore profile of decreasing wind speed toward the coast), current feedback on the surface stress (causing a large dampening of the mesoscale activity; Renault et al. (2016d, 2019a)), and high-frequency wind fluctuations.

In this paper and its biogeochemical companion (Deutsch et al., 2021a), ROMS is implemented over the CCS and is forced by the atmosphere with a regional configuration of the Weather Research Forecast (WRF) model for the period 1995-2010. The main objectives are to characterize and validate the behavior of the CCS circulation at different time scales with good mesoscale resolution in both the ocean (*dx* = 4 km) and atmosphere (*dx* = 6 km), while also reviewing the now substantial literature on this relatively well measured regional system. We also also provide an assessment of the importance of synoptic wind forcing on the mean and mesoscale currents as well as on the inertial currents. Overall, it provides a more comprehensive validation assessment than is customary, both to establish the credentials of this particular model for its intended applications (mostly biogeochemical and ecological, *e.g*., Deutsch et al. (2021a)) and to provide an example of the state of the art for realistic regional simulations.

In our view, “realistic” model simulations — using forcing and bathymetry fields derived from measurements and parameterizations for the subgrid-scale effects perceived to be essential — are coming to play an increasingly central role in oceanic sciences. It is therefore important to develop a better sense in the community of just how accurate such a virtual reality is, as well as what its limitations are (*e.g*., McWilliams, 2007). This is a necessary maturation step for this oceanic methodology, as it has long since been one for global climate science.

The datasets and the model components, setup, and analysis methodology are described in Sec. 2. In Sec. 3, the behavior of the atmospheric forcing is evaluated with respect to satellite measurements. Section 4 aims to evaluate the oceanic circulation and subsurface layer using satellite and *in situ* measurements. Finally, in Sec. 5, the oceanic mesoscale activity is evaluated and the importance of the high frequency atmospheric forcing is assessed. The results are discussed and summarized in Sec. 6.

## 2 Model Configurations, Analysis Methods, and Data

### 2.1 The Regional Oceanic Modeling System (ROMS)

The oceanic simulations are made with ROMS (Shchepetkin and McWilliams, 2005; Shchepetkin, 2015). As in Renault et al. (2016d), the primary U. S. West Coast (USW4) simulation domain extends from 144.7°W to 112.5°W and from 22.7°N to 51.1°N. Its horizontal grid is 437 × 662 points with a resolution of *dx* = 4 km, and it has 60 terrain-and surface-following sigma levels in the vertical with stretching parameters *h_cline_* = 250 m, and *θ_b_* = 3.0, and *θ_s_* = 6 (Shchepetkin and McWilliams, 2009).

Initial and horizontal boundary data for *T, S*, surface elevation, and horizontal velocity are taken from the quarter-degree, daily-averaged Mercator Glorys2V3 product (http://www.myocean.eu), and applied to the outer boundary of a *dx* = 12 km solution, which spans a larger domain and serves as a parent grid for the USW4 solution. To improve the water mass representation, in particular the density distribution, the Mercator data are corrected using the mean monthly climatology from the World Ocean Atlas (WOA) (Locarnini et al., 2013; Zweng et al., 2013) over the period 1995-2004. As we shall see in Sec. 4.2, the model does exhibit a mean bias in *S* (*e.g*., the geographical distribution on an interior density surface), and our understanding is that this is mostly inherited from WOA due to the sparsity of interior hydrographic measurements used to determine an accurate mean state around the model boundaries.^2^

The surface turbulent evaporation, heat, and momentum fluxes are estimated using bulk formulae (Large, 2006), and the atmospheric surface fields are derived from an uncoupled WRF simulation (Sec. 2.2), along with the precipitation and downwelling radiation; for these surface fluxes the temporal sampling interval is one hour (1H; see Sec. 5.2 for the sensitivity to this interval). As in Lemarié et al. (2012), the river-runoff forcing dataset we use is a monthly climatology from Dai et al. (2009). River runoff is included offline as surface precipitation and is spread using a Gaussian distribution over the grid cells that fall within the range from the coast to 150 km offshore; this excludes a detailed representation of river plumes.

When forced with bulk formulae, uncoupled oceanic simulations often estimate the surface stress using the absolute wind vector ***U**_a_* (*e.g*., at 10 m height). As shown by *e.g*., Dewar and Flierl (1987); Duhaut and Straub (2006); Eden and Dietze (2009); Renault et al. (2016d,c); Jullien et al. (2020), such simulations overestimate the mesoscale activity because of their lack of a current feedback. The current feedback is simply the influence of the surface current on the surface stress and low-level wind. In a coupled ocean-atmosphere model, the relative velocity difference between the surface wind and current ***U**_r_* is used in the bulk formula, ***τ*** = *ρ_a_C_d_* |***U**_r_* | ***U**_r_* (with *ρ_a_* the surface air density and *C_d_* the drag coefficient). Although the 10-m is generally much larger than the surface current (*e.g*., when *U_a_* = 10 ms ^−1^ and *U_o_* = 1 m s ^−1^, *U_r_* = 9 m s ^−1^), at the mesoscale the current feedback induces a sink of energy from the currents to the atmosphere, which causes a large dampening of the mesoscale activity (by ≈ 40% for the U. S. West Coast; Seo et al. (2016); Renault et al. (2016d)). However, in a forced oceanic model, an opposite bias arises: the mesoscale activity is underestimated because the wind response to the weakened currents through this stress feedback should partially re-energize the atmosphere, hence also the mesoscale currents. Renault et al. (2016d, 2020) suggest using a wind-correction approach based on the current-wind coupling coefficient *s_w_*, estimated from a coupled simulation as the slope between the mesoscale current vorticity and the mesoscale surface stress curl. The atmospheric re-energization is then expressed as

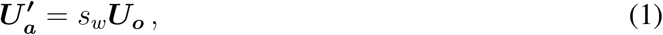

where ***U**_o_* is the surface current, 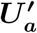 is the wind response to ***U**_o_*, and *s_w_* is a statistical regression coefficient. The surface stress, therefore, is computed using a bulk drag formula,

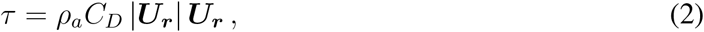

with a parameterized relative velocity, ***U**_r_*:

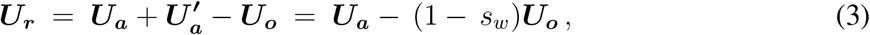

where ***U**_a_* is the surface wind from an uncoupled atmospheric product. For the CCS region, *s_w_* = 0.23 ± 0.1 (Renault et al., 2016d, 2019b). For instance, a surface current of 1 m s^−1^ is expected to induce a 10-m wind anomaly of 0.23 ms^−1^. This simple parameterization roughly mimics the wind response to the current feedback. Although such a parameterization presents some limitations (*i.e*., in this study *s_w_* is constant, so does not take into account the seasonal cycle, the atmospheric boundary layer dependency, nor the global-scale geographic variation of *s_w_*), but it does lead to approximately the expected dampening and re-energization of the mesoscale currents.

The statistically equilibrated solution USW4 is integrated over the period 1995-2010 after a spin up of 1 year starting from a larger-domain ROMS parent solution with *dx* = 12 km.

### 2.2 The Weather Research and Forecast Model (WRF)

WRF (version 3.6.1; Skamarock et al. (2008)) is implemented in a configuration with two grids, similar to Renault et al. (2016b). The WRF domains are slightly larger than the ROMS domains to avoid the effect of the WRF boundary sponge (4 grid points wide). It has horizontal resolutions of *dx* = 18 km and 6 km, respectively, using only the latter over the USW4 domain. The model is initialized with the Climate Forecast System Reanalysis (CFSR with *dx* ≈ 40 km horizontal resolution; Saha et al. (2010)) from 1 January 1994 and integrated for 17 years with time-dependent boundary conditions interpolated from the same six-hourly reanalysis. Forty vertical levels are used, with half of them in the lowest 1.5 km, as in Renault et al. (2016b). The model configuration is set up with the same parameterizations as in Renault et al. (2016b) except that the WRF Single-Moment, 6-class microphysics scheme (Hong and Lim, 2006) is modified to take into account the spatial and seasonal variations of the droplet concentration (Jousse et al., 2016). Its Sea Surface Temperature (SST) forcing is derived from the Ostia one-day product (Stark et al., 2007) that has a spatial resolution of *dx* = 5 km. The inner-nested domain (WRF6) is initialized from the outer solution (WRF18) on 1 April 1994 and integrated for 17 years. Only the period 1995-2010 is used in the model evaluations.

### 2.3 Analysis Methods

The numerical outputs for the solutions are daily averages, except when assessing the high frequency forcing importance where hourly averages outputs are saved. The winter, spring, sum-mer, and fall seasons correspond to the months January-March, April-June, July-September, and October-December, respectively. To assess the realism of the oceanic and atmospheric solutions, several sub-regions are considered (Fig. 1): see the separate boxes for southern California (South), central California (Central), and northern California plus Oregon and Washington (North). Additionally, when the the data have a spatial resolution that is high enough to consider a coastal region (i.e., *dx* < 1°) and do not have too large a nearshore bias zone (e.g., QuikSCAT products usually have a coastal blind zone about 30-50 km wide), both Nearshore and Offshore sub-boxes are also considered. The mean (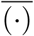) is defined with respect to the full time average (1995-2010), and it is done separately for each season; the prime (·)′ denotes a deviation from the mean.

**Figure 1:**
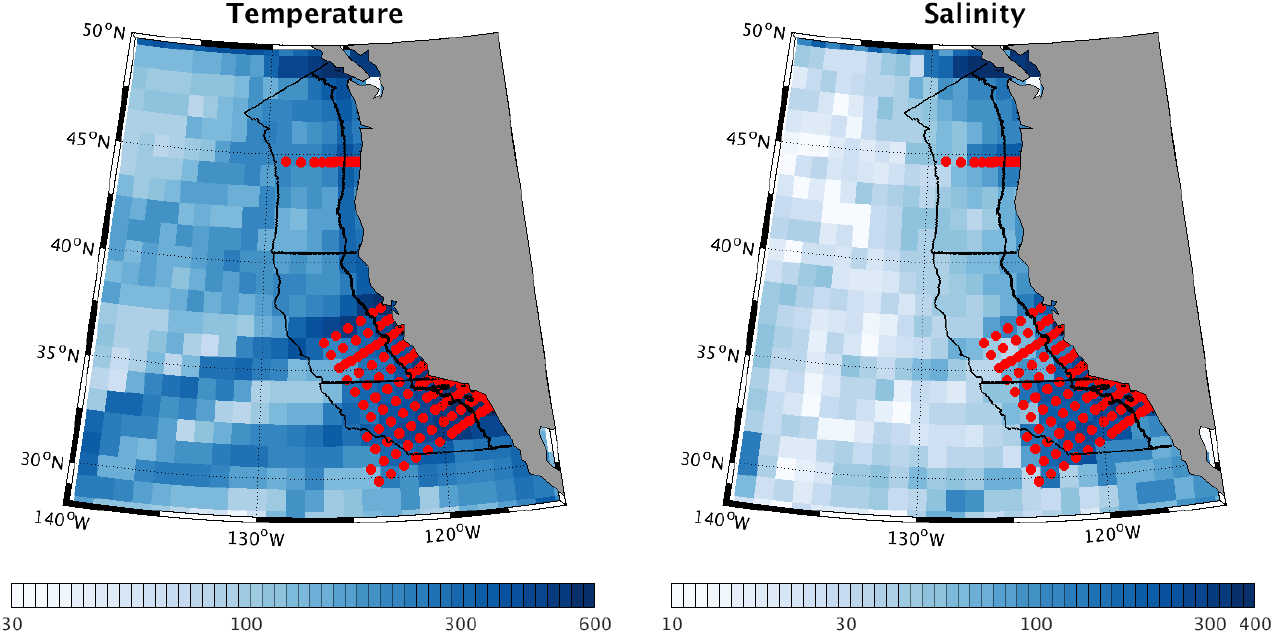
Data density of temperature and salinity measurements in World Ocean Database. The black lines indicate the box areas used to evaluate the simulations with respect to the measurements. Three alongshore domains are assessed: South, Central, and North. When the data have a high-enough spatial resolution to assess a coastal region (*i.e*., more than two cross-shore valid data), Nearshore and Offshore boxes are also considered with the meridional boundary line, also in black. The red circles indicate the CalCOFI stations and the Newport hydrographic line.

The oceanic geostrophic surface currents are estimated using daily-averaged sea surface height:

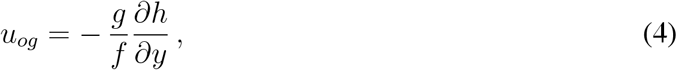

and

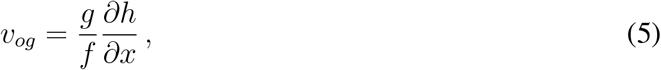

where *u_og_* and *v_og_* are zonal and meridional geostrophic currents, *g* is gravitational acceleration, *f* is Coriolis frequency, and *h* is sea surface height.

Following the method described in Renault et al. (2016c), the total wind work is defined as

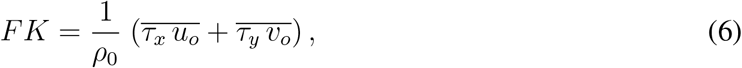

where u_o_ and *v_o_* are the zonal and meridional surface currents, *τ_x_* and *τ_y_* are the zonal and meridional surface stresses, and *ρ_0_* is the mean seawater density. Substituting the decomposition of (4) and (5) into (6), the total wind work on the geostrophic and ageostrophic flow are

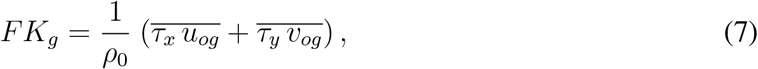

and

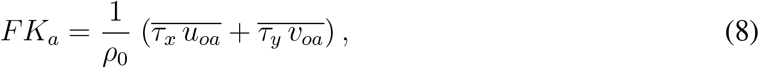

where *u_oa_* and *v_oa_* are the zonal and meridional components of ageostrophic velocities, respectively. The wind work terms *FK_g_* and *FK_a_* can be split into their mean (*F_m_K_mg_* and *F_m_K_ma_*) and eddy parts (*F_e_K_eg_* and *F_e_K_ea_*):

- mean geostrophic wind work,

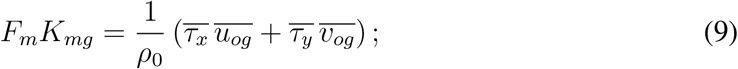
- mean ageostrophic wind work,

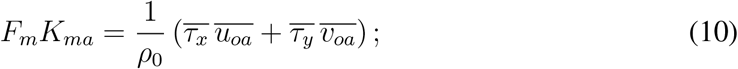
- geostrophic eddy wind work,

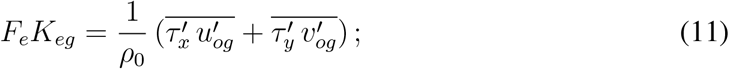
- ageostrophic eddy wind work,

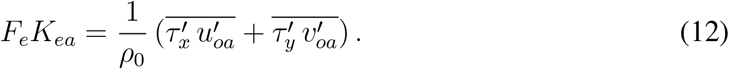 As in Stern (1975), Marchesiello et al. (2003), and Renault et al. (2016d), we evaluate the following relevant eddy-mean energy conversion terms:
- barotropic (horizontal Reynolds stress) kinetic energy conversion *K_m_K_e_*,

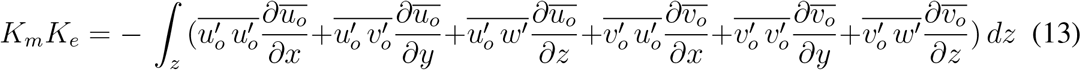

(where *w* is the vertical velocity and x, *y*, and *z* are the zonal, meridional, and vertical coordinates, respectively) and
- eddy baroclinic potential-to-kinetic conversion *P_e_K_e_*,

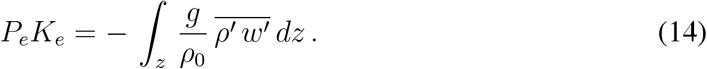

*F_m_K_mg_* represents the transfer of energy from mean surface wind forcing to mean geostrophic kinetic energy; *F_m_K_ma_* represents the transfer of energy from mean surface wind forcing to mean ageostrophic kinetic energy; *F_e_K_eg_* represents the transfer of energy from surface wind forcing anomalies to geostrophic EKE; *F_e_K_ea_* represents the transfer of energy from surface wind forcing anomalies to ageostrophic EKE; *K_m_K_e_* represents the barotropic conversion from mean kinetic energy to EKE; and *P_e_K_e_* represents the baroclinic conversion from eddy available potential energy to EKE. We compute those conversion terms at each model grid point. The wind work is estimated at the free surface, while the barotropic and baroclinic conversion terms are integrated over the whole water column.

### 2.4 Primary Observational Datasets

Satellite and in *situ* measurements are used to evaluate the realism of both the atmospheric and oceanic simulations. Because of intermittent sampling with different instruments, we do not insist on exact time correspondences in computing climatological averages. To evaluate the performance of the atmospheric simulation in terms of cloud cover, we use remote sensing data retrieved from the Moderate Resolution Imaging Spectrometer level 2 data (MODIS; Platnick et al. (2003)). We use data from the Terra satellite, which is available twice daily around 10:30 am/pm local time, beginning in the year 2000. The Forcing for Coordinated Ocean-ice Reference Experiments 2 (CORE; Large and Yeager (2009)) dataset is used to evaluate the surface heat and freshwater fluxes. It provides monthly surface fluxes at a spatial resolution of 1°. The monthly Global Precipitation Climatology Project (GPCP; Adler et al. (2003)) is also used to evaluate precipitation. It has a spatial resolution of 1°. The surface stress data is from the Scatterometer Climatology of Ocean Winds (SCOW; Risien and Chelton (2008)) product based on the QuikSCAT satellite scatterometer. It provides monthly data at a 0.25°resolution. To compute the wind work, the surface stress daily product processed by the Centre ERS d’Archivage et de Traitement (CERSAT; Bentamy and Fillon (2012)) is used. It provides daily surface stress at a spatial resolution of *dx* = 25 km. SST forcing is derived from the Ostia daily product (Stark et al., 2007) that has a spatial resolution of *dx* = 5 km. We use data from the California Cooperative Oceanic Fisheries Investigations (CalCOFI) (e.g., Bograd et al. (2003)). Since 1950 hydrographic stations have been repeatedly but irregularly sampled on a geographically fixed grid. In this study line 80 (off Pt. Conception; 34°N) is used to estimate a seasonal climatology of temperature, salinity, and density, respectively, to validate the simulation from 1995 to 2010. The temperature, salinity, and density are further evaluated using the World Ocean Database 2013 (WOD13, Locarnini et al. (2013); Zweng et al. (2013)). Its fields have a resolution of *dx* = 25 km and extend from the surface to the bottom of the ocean. The WOD13 dataset for that period includes the CalCOFI data and the Newport hydrographic line. The CSIRO (Commonwealth Scientific and Industrial Research Organization) Atlas of Regional Seas (CARS) climatology (Ridgway et al., 2002) provides an estimate of the monthly climatology of the Mixed Layer Depth (MLD) using a temperature threshold of ΔΘ = 0.2 °and Δσθ = 0.03 kg m^−3^. Finally, the CNES-CLS13 dataset (Rio et al., 2014) is used to evaluate the simulated mean sea surface height and to estimate the geostrophic wind work. It is a combination of GRACE satellite data, altimetry, and *in situ* measurements with a spatial resolution of *dx* = 25 km in the analysis product. The Archiving, Validation, and Interpretation of Satellite Oceanographic Data (AVISO) dataset (Ducet et al., 2000) is used to evaluate the mesoscale activity simulated by USW4 and to estimate geostrophic wind work. It provides the daily sea level anomaly at a resolution of *dx* = 25 km. Finally, the gliders (line 66.7) of Rudnick et al. (2017) are used to evaluate the geostrophic current structure.

## 3 Atmospheric Fields

### 3.1 Shortwave Radiation

Surface net shortwave flux is a key component of the surface energy budget in the CCS. In numerical models, it is strongly related to the representation of clouds and their radiative properties, which is a common difficulty for both global and regional climate models in eastern boundary upwelling regions (Nam et al., 2012; Wyant et al., 2010; Zermeño-Díaz et al., 2015). The difficulty is at least partially attributable to approximate parameterizations of processes governing the stratocumulus clouds that dominate these regions due to the combination of large-scale tropospheric downwelling, low humidity, and a cold oceanic SST adjacent to a generally warmer continent. WRF offers choices among various physical parameterizations; here we make our choice in accordance with previous work where an optimized combination of parameterizations was established in WRF for a stratocumulus region (Jousse et al., 2016). In particular, this combination (Sec. 2.2) minimize stratocumulus biases, and for the present WRF simulations similar sensitivity tests confirm the previous choices. We also perform sensitivity tests for the surface shortwave radiation scheme (not done in Jousse et al. (2016)). Our results show a better performance of the Goddard Shortwave scheme (Chou and Suarez, 1994) in comparison to the Dudhia scheme (Dudhia, 1989). We prescribe the observed spatial variability and seasonality for the cloud droplet concentration number in the microphysics parameterization scheme WSM6 Jousse et al. (2016). This modeling strategy minimizes biases in the liquid water path over the northeast Pacific. Figure 2 demonstrates the plausible WRF results for both spatial variability and seasonal cycle in all the regions of interests. There is a general increase in incident flux moving equatorward and, in the south, shoreward, modulated by the cloud cover. These results reflect the realism of the cloud macrophysical structure (*i.e*., total water path, TWP) in the simulation (Jousse et al., 2016). Along the central California coast, both cloud cover and mean shortwave fluxes are biased with respect to the measurements. While no doubt some of these are due to model errors, near the coast the satellite measurements have a too coarse spatial resolution to resolve the nearshore variability. There is also an underestimation of the shortwave flux over the Southern California Bight caused by the over-estimation of the cloud cover by 5-10 % (not shown). These biases are relatively small compared to global climate models (more than 30%, see *e.g*., Fig. 2 of Richter (2015)). The behavior of the model in reproducing the interannual variability of the shortwave radiation is also revealed in Fig. 3abc. It depicts the interannual variation of the yearly mean net shortwave radiation averaged over the South, Central, and North Boxes for CORE and USW4. The simulation fairly reproduces the interannual variability with *e.g*., a more intense shortwave radiation in 1997 (+10 W m^−2^) with respect to the other years.

**Figure 2:**
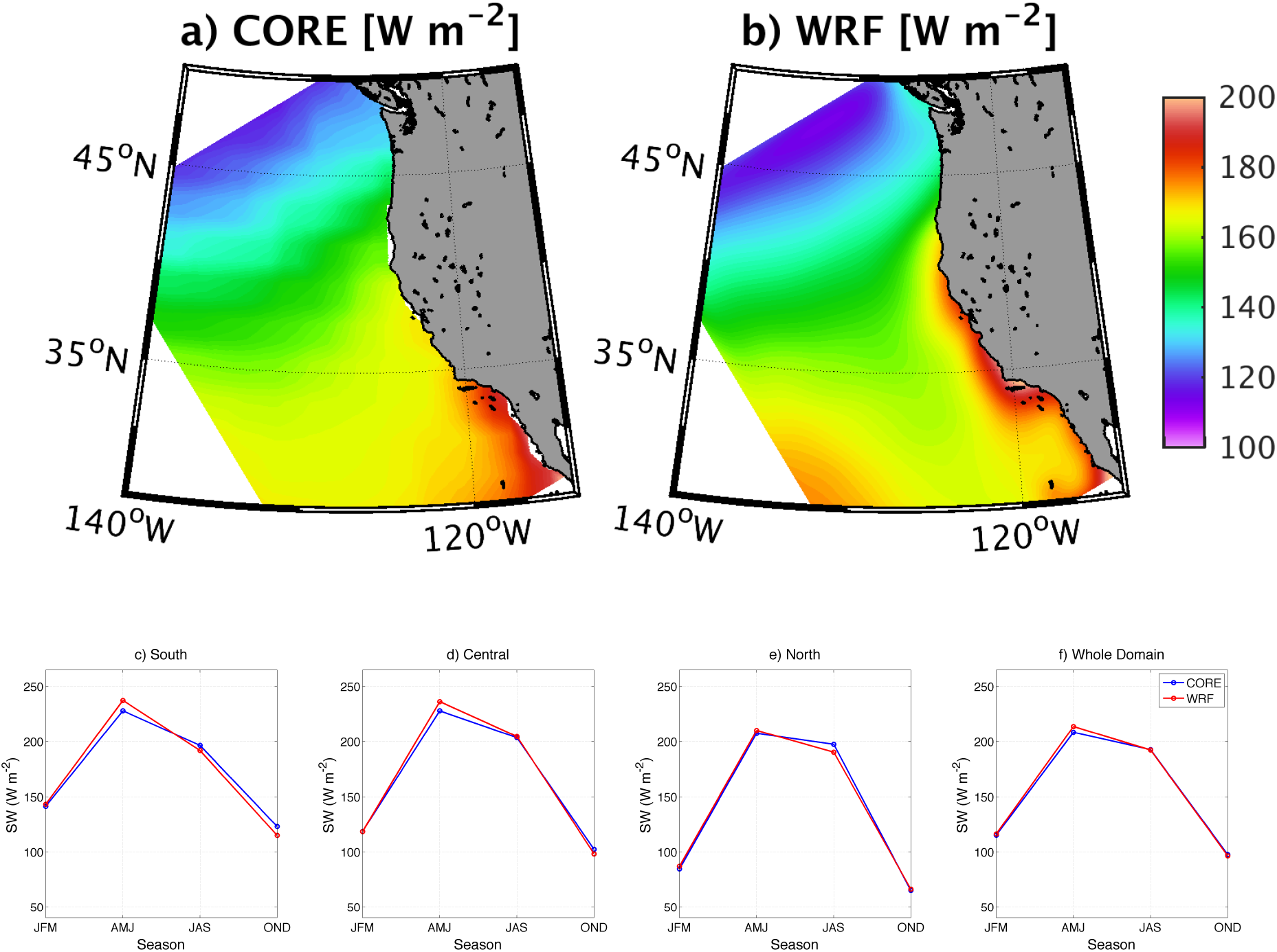
Mean shortwave radiation [W m^−2^] estimated for the period 1995-2006 from (a) CORE and (b) USW4. Panels (c), (d), (e), and (f) represent the seasonal shortwave radiation variation estimated over the same period from CORE (blue) and WRF (red), averaged over the boxes indicated in Fig. 1 or over the whole domain. The realistic representation of the cloud cover and of the liquid water path in the model allows a good representation of the shortwave radiation.

**Figure 3:**
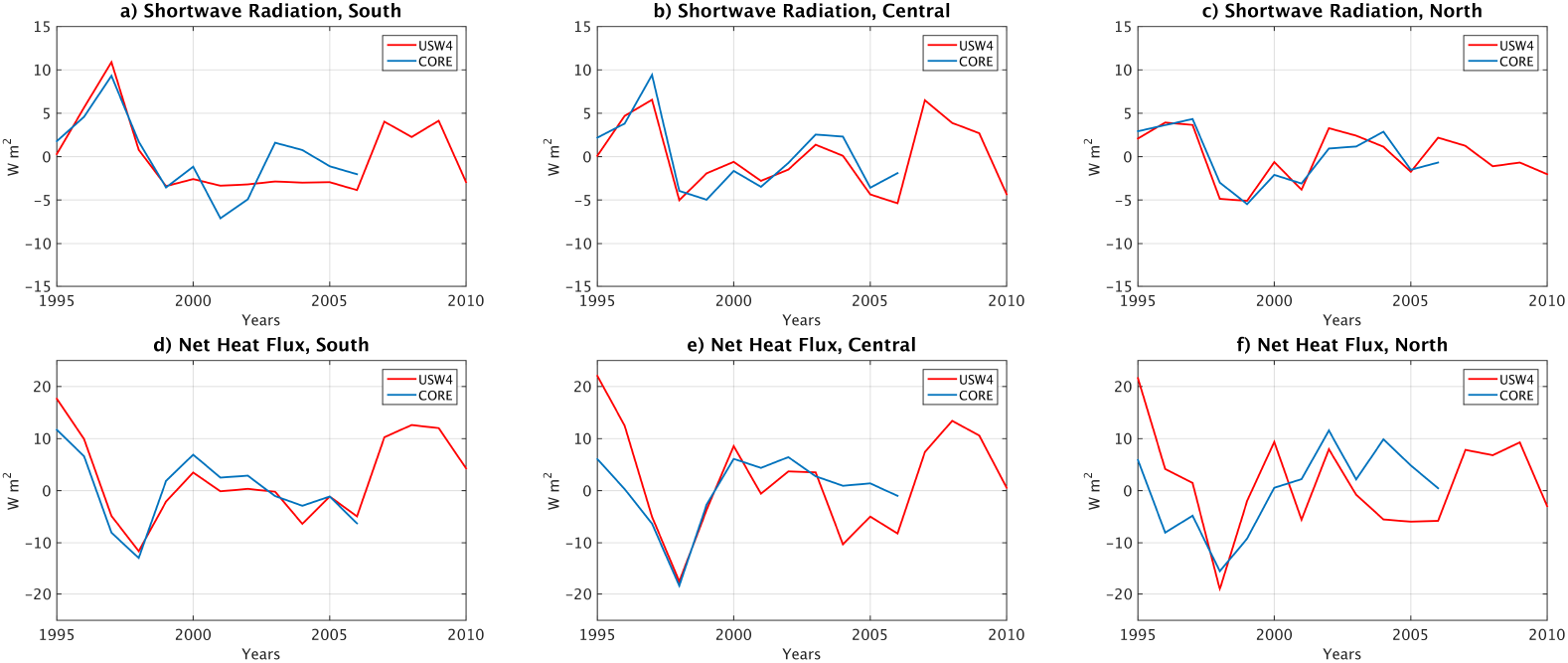
Interannual shortwave radiation (abc) and net radiation (def) [W m ^−2^] over the boxes indicated in Fig. 1 from CORE and USW4.

### 3.2 Surface Heat Flux

The net surface heat flux is calculated in USW4 using the WRF solar radiation, 10 m wind, 2 m temperature and humidity fields, and SST. The resulting flux is evaluated with respect to the CORE dataset. Figure 4 shows the mean net surface heat flux and its seasonal variability in the different regions (i.e., the alongshore boxes in Fig. 1). Here, we do not consider Offshore and Nearshore boxes because of the coarse spatial resolution of CORE (1°). Despite some bias compensations, there is an overall good agreement between the measurements and the simulations both in terms of spatial variability and seasonal cycle. Due to the upwelling and the cold coastal SST in the CCS, there is generally strong net heating of the ocean near the coast. The maximum of net heat flux in the Central coast box is similar in both the observations and USW4. Consistent with the overestimation of the cloud cover and the underestimation of the shortwave radiation, during the upwelling seasons (spring and summer), the largest bias is again located in the Southern California Bight, where the net surface heat flux is underestimated by 10 W m^−2^. Along the central California coast, the cloud cover bias induces a positive bias in shortwave radiation that is mostly compensated for by a negative bias in longwave radiation (not shown). The turbulent (latent and sensible) heat fluxes exhibit less than a 7% error (*i.e*., too large a latent heat flux), which is within the error range for the measurements (Large and Yeager, 2009). Finally, near the coast in a band ≈ 30 km wide, the net heat flux is higher than in the measurements, which probably reflects the coarser spatial resolution of CORE. The realistic representation of the net heat flux leads to a reasonably good estimate of the spatial and the seasonal variation of the SST (Sec. 4.1); however, as shown in the companion paper (Deutsch et al., 2021a), the shortwave flux bias along the California coast can induce a positive bias in chlorophyll through photosynthesis. Figure 3def depicts the interannual yearly variation of the net heat fluxes over the South, Central, and North boxes for CORE and USW4. Again, the interannual variability is fairly reproduced by the simulation. Interestingly, the peak of shortwave radiation in 1997 is compensated by more intense turbulent heat fluxes (not shown).

**Figure 4:**
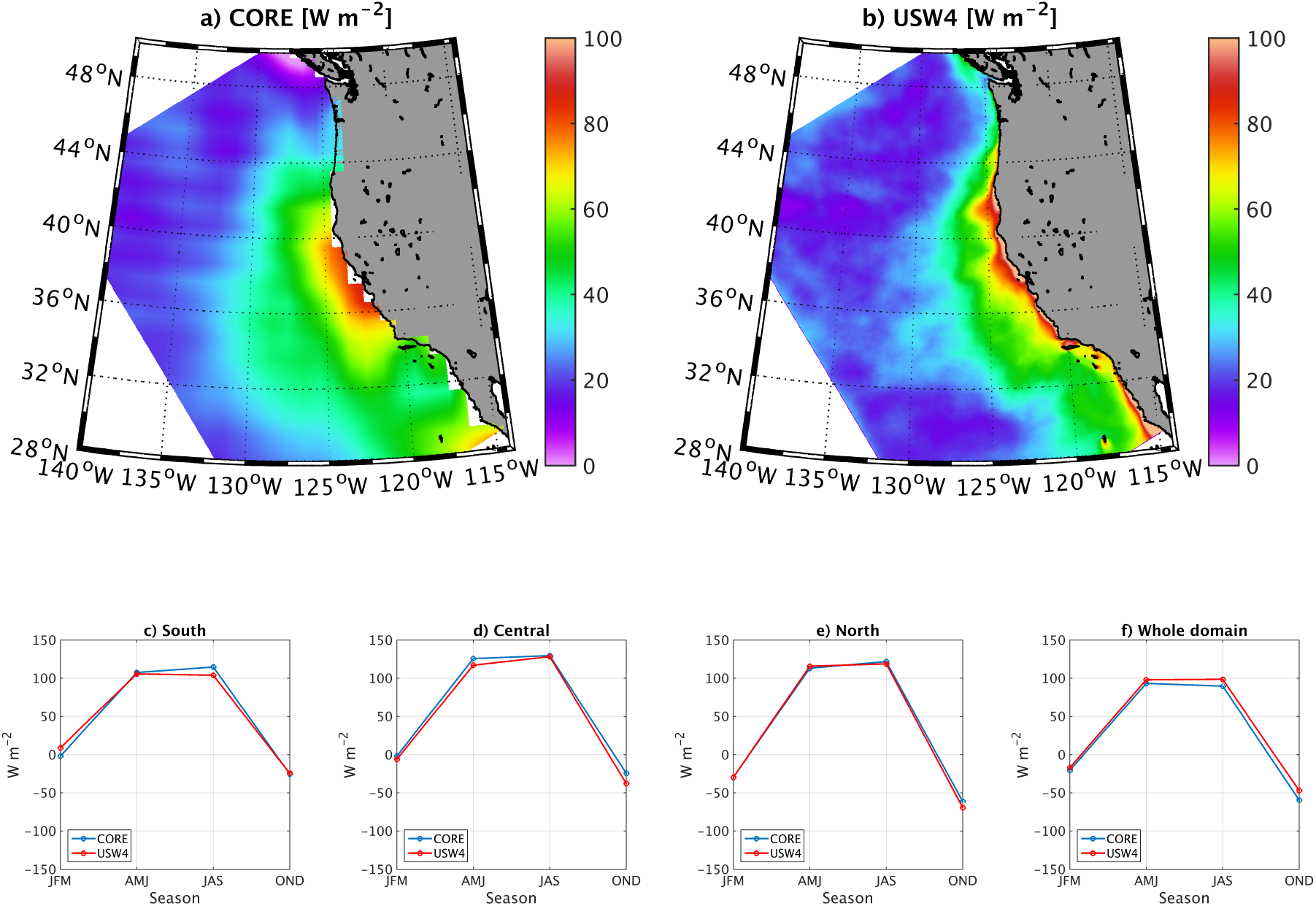
Mean net heat fluxes [W m^−2^] from (a) CORE and (b) USW4 over the period 1995-2006. Panels (c), (d), (e), and (f) represent the seasonal evolution of the net heat fluxes over the same period from CORE (blue) and USW4 (red), averaged over the boxes indicated in Fig. 1 or over the whole domain.

### 3.3 Surface Freshwater Flux

The net freshwater flux (evaporation minus precipitation) is computed by combining the precipitation from WRF and the evaporation estimated by bulk formulae from WRF’s surface fields (Large, 2006). Evaporation dominates toward the south over the warmer subtropical gyre, and precipitation dominates to the north, especially close to the coast; *i.e*., there is more precipitation in the north during the winter months due to the storm tracks. WRF generally overestimates GPCP by about 0.5 to 1 mm day^−1^ (not shown). These differences are certainly not negligible values for the water budget. Nevertheless, they remain within the observational uncertainty range provided by the GPCP data. Moreover, due to their lack of sensitivity to drizzle, remote sensors are known to generally underestimate the precipitation produced by low clouds (Rapp et al., 2013). Because the CCS is substantially covered with low clouds, this may explain some of the discrepancies between WRF and GPCP (*n.b*., similar results are found using CORE). In USW4, the overestimation of the precipitation is compensated by a slight excess of evaporation (consistent with the latent heat flux bias), which leads to a realistic agreement of the net freshwater flux between USW4 and CORE (Fig. 5). In both USW4 and the observations, the net freshwater flux does not have a strong inter-annual variability as shown in Fig. 6abc except over the South Box, where it reaches large values in 2009 and 2010.

**Figure 5:**
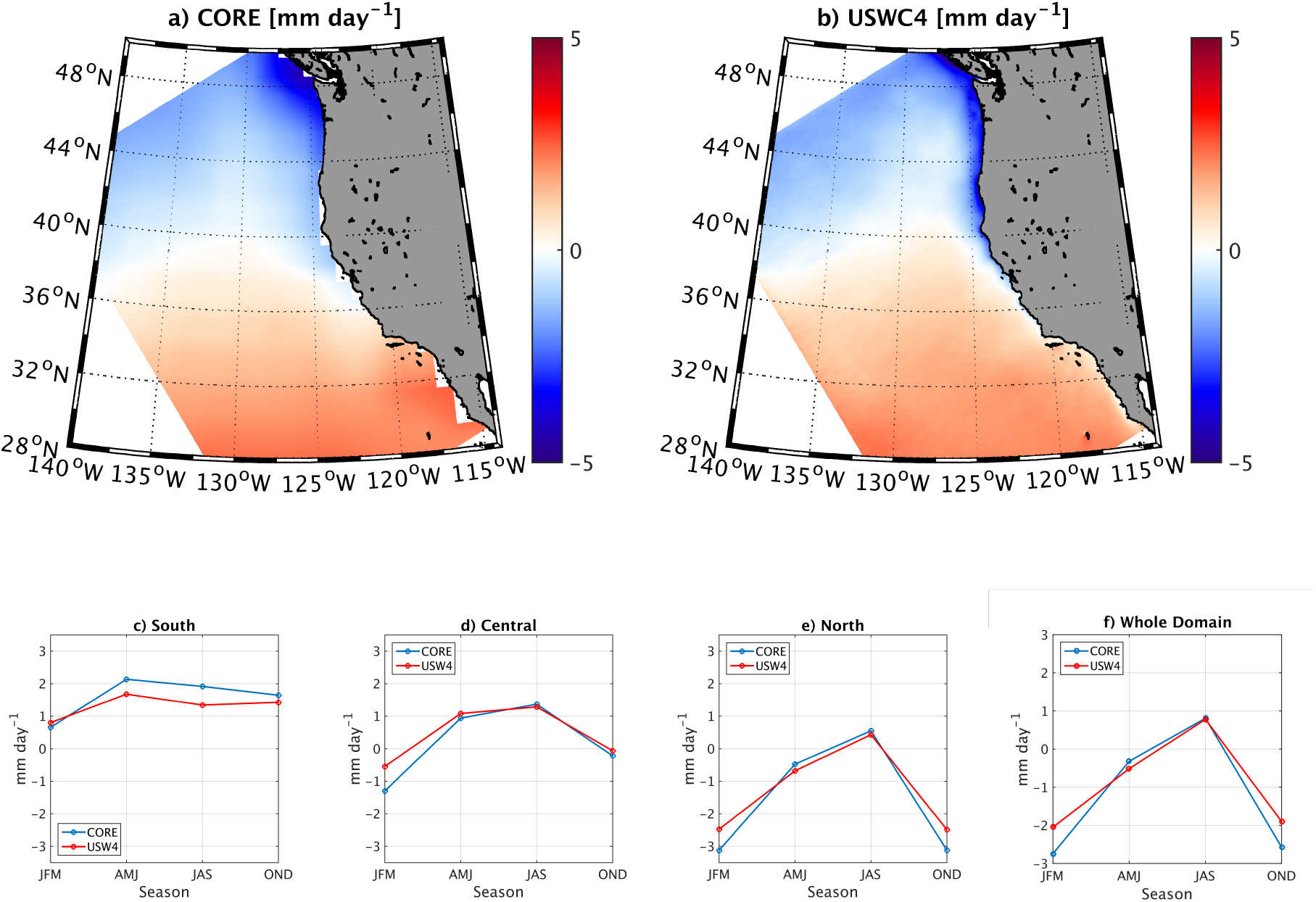
Net surface freshwater flux [mm day^−1^] from (a) CORE and (b) USW4 over the period 1995-2006. Panels (c), (d), (e), and (f) represent the seasonal evolution of the net surface freshwater flux over the same period from CORE (blue) and USW4 (red), averaged over the boxes indicated in Fig. 1 or over the whole domain. The simulation reproduces the main spatial pattern of the observed freshwater flux and its seasonal variability.

**Figure 6:**
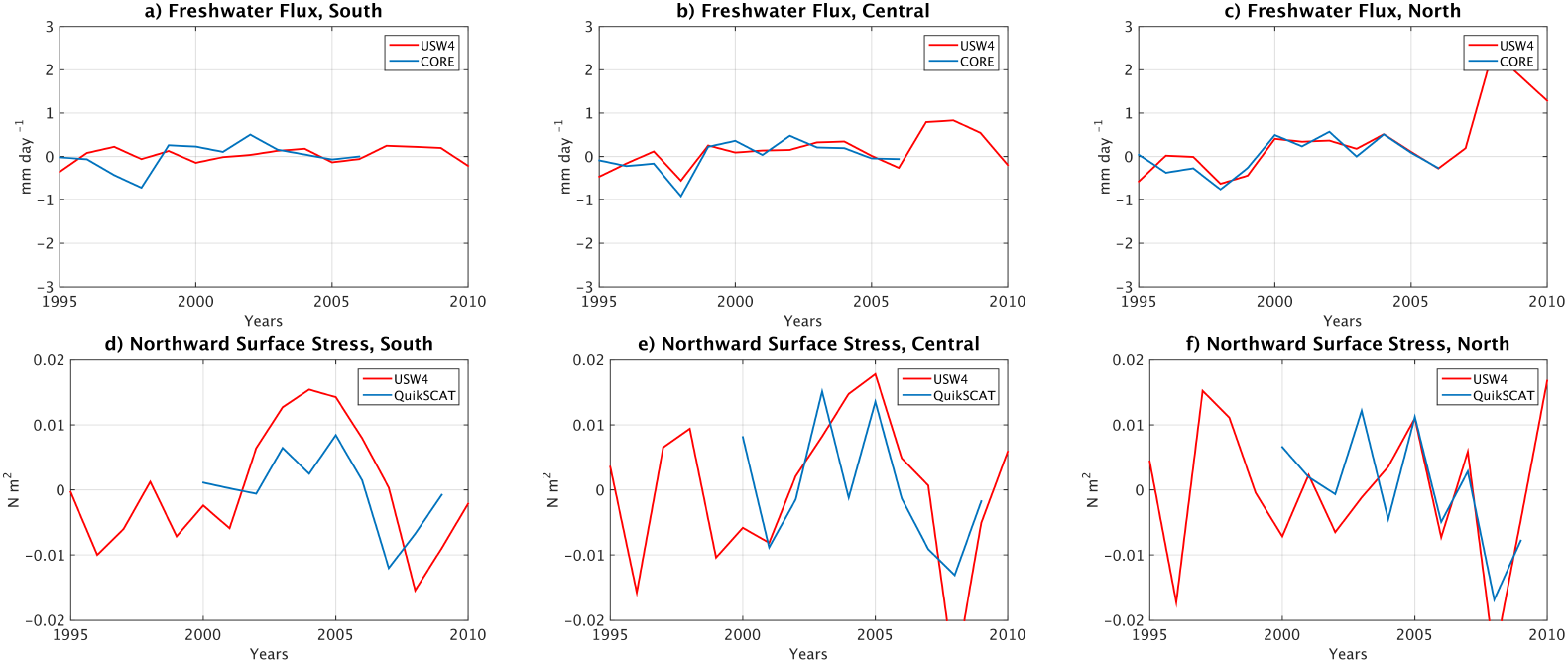
Same as Fig. 3 but for the net freshwater flux [mm day ^1^] from CORE and northward surface stress [N m^−2^] from QuikSCAT.

### 3.4 Surface Stress

The surface stress is calculated in USW4 with the Large (2006) bulk formulae as described in Sec. 2.1; the 10 m wind, and the 2 m temperature and humidity. Renault et al. (2016b) show a good agreement between the 10 m wind and satellite and *in situ* observations with a similar model configuration. Here, we evaluate the simulated surface stress with respect to SCOW (Fig. 7). In both USW4 and SCOW, over the CCS the surface stress is mostly equatorward due to the offshore position of the atmospheric subtropical high, and this is the primary cause of offshore Ekman transport and coastal upwelling. This pattern is persistent in the South and Central boxes, with peak stresses near the coast in spring and summer. In the north, the alongshore wind stress direction reverses seasonally. The simulated surface stress is similar to that in the observations in both amplitude and direction. The seasonality and the main gradients are also realistic (Fig. 7). We do not consider separate Nearshore and Offshore boxes because, as noted by *e.g*., Renault et al. (2009, 2016b), QuikSCAT data do not measure the stress within the first ≈ 30 km from the coast due to land contamination in the backscatter measurements (Chelton et al., 2004). The upwelling season in spring and summer is marked by a distinctive alongshore surface stress (up to 0.09 N m^−2^), and the numerous capes and mountain ranges induce so-called expansion fans (Winant et al., 1988). The main discrepancies between SCOW and WRF occur close to the coast. In the simulation there is a coastal band where the surface stress is reduced compared to its offshore value (*i.e*., the wind drop-off). Such a slackening of the wind is mainly caused by the presence of coastal orography, coastline shape, the difference between marine and terrestrial drag coefficients, and SST; this drop-off pattern is not well captured in QuikSCAT. An indirect validation of the wind drop-off is given by the oceanic response. A too wide drop-off causes a poor representation of mean oceanic current structure and mesoscale activity (Renault et al., 2016b). Finally, the stress magnitude is slightly underestimated by USW4: the mean biases over the whole domain are 0.006 N m^−2^ and 0.003 N m^−2^ for the meridional and the zonal surface stress, respectively, *i.e*., the same order of magnitude as the QuikSCAT surface stress error (Risien and Chelton, 2008). The least skillfully simulated region is again the Southern California Bight, where the surface stress is weak compared to other parts of the CCS, and it is overestimated in USW4; perhaps the complicated small island geometry is a contributing cause. Similar results are found using the QuikSCAT interannual product (Bentamy and Fillon, 2012).

**Figure 7:**
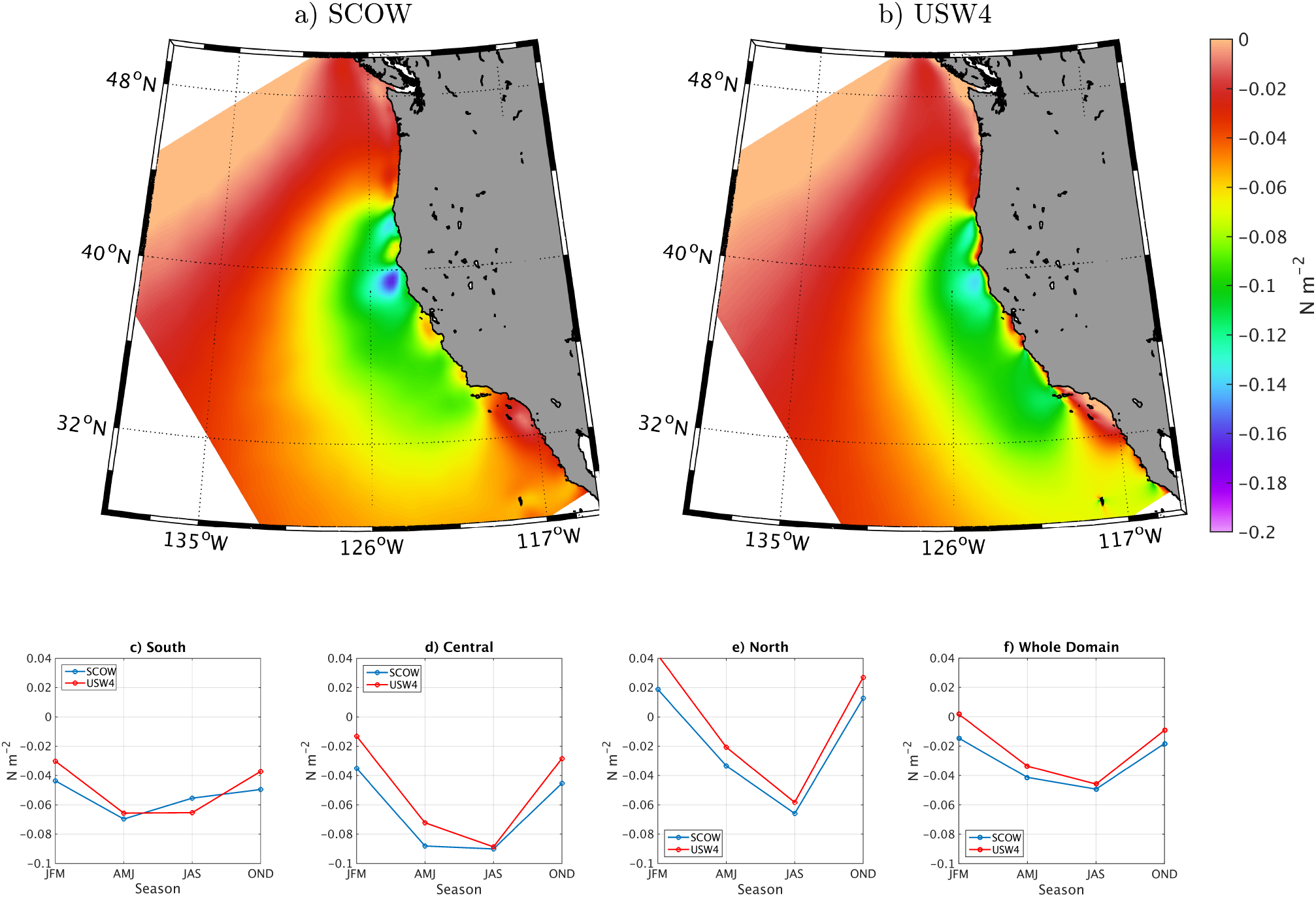
Mean meridional surface stress from (a) SCOW and (b) USW4 for the upwelling season (spring and summer) estimated over the period 2000-2009. Panels (c), (d), (e), and (f) represent the seasonal evolution over the same period from SCOW (blue) and USW4 (red), averaged over the boxes indicated in Fig. 1 or over the whole domain. USW4 reproduces the main surface stress spatial pattern and its seasonal evolution.

The surface stress curl during the upwelling season from SCOW and USW4 is then compared in Fig. 8 to CFSR and NAM that are widely used to force oceanic models. As noted previously, because of the blind zone of QuikSCAT, SCOW does not represent nearshore large positive values of the surface curl caused by the wind drop-off. USW4 has large positive values of surface stress curl. Consistent with Renault et al. (2016b), the surface stress curl has spatial and seasonal variability (not shown) in both its offshore extent and intensity. The offshore extent varies from around 10 to 80 km from the coast and the surface stress reduction from 10 to 80 % (corresponding to the largest values of the curl). The largest curl values are situated when the mountain orography is combined with the coastline shape of a cape. Interestingly, the Santa Barbara Channel is characterized by the presence of many fine-scale wind structures that are induced by cape effects and island mass effects. As an indirect validation, Kessouri et al. (2021a), shows that the these fine-scale wind structures are responsible for intense blooms. The surface curl from CFSR does represent a drop-off, however, it is characterized by a too large cross-shore extent that is a characteristic of too coarse spatial resolution atmospheric model (≈ 35 *km*). It also presents a pattern of positive, negative and positive values parallel to the coast, which is characteristic of the Gibbs phenomenon in spectral models (Hoskins, 1980). CFSR does not either represent any of the fine-scale wind structure along the coast. As shown by Renault et al. (2016a) and Kessouri et al. (2021a), such a wind product does not allow to represent the current structure nor the mesoscale activity and, thus, the net primary production along the CCS. NAM is a reanalysis over the United States of America from NCEP. It is a configuration of the WRF model with a spatial resolution of *dx* = 12 km. Although the representation of the wind drop-off is improved with respect to CFSR, the spatial extent of the drop-off is still too large and it does not represent all the fine-scale structures simulated by our simulations such as the island mass effects over the Santa Barbara Channel.

**Figure 8:**
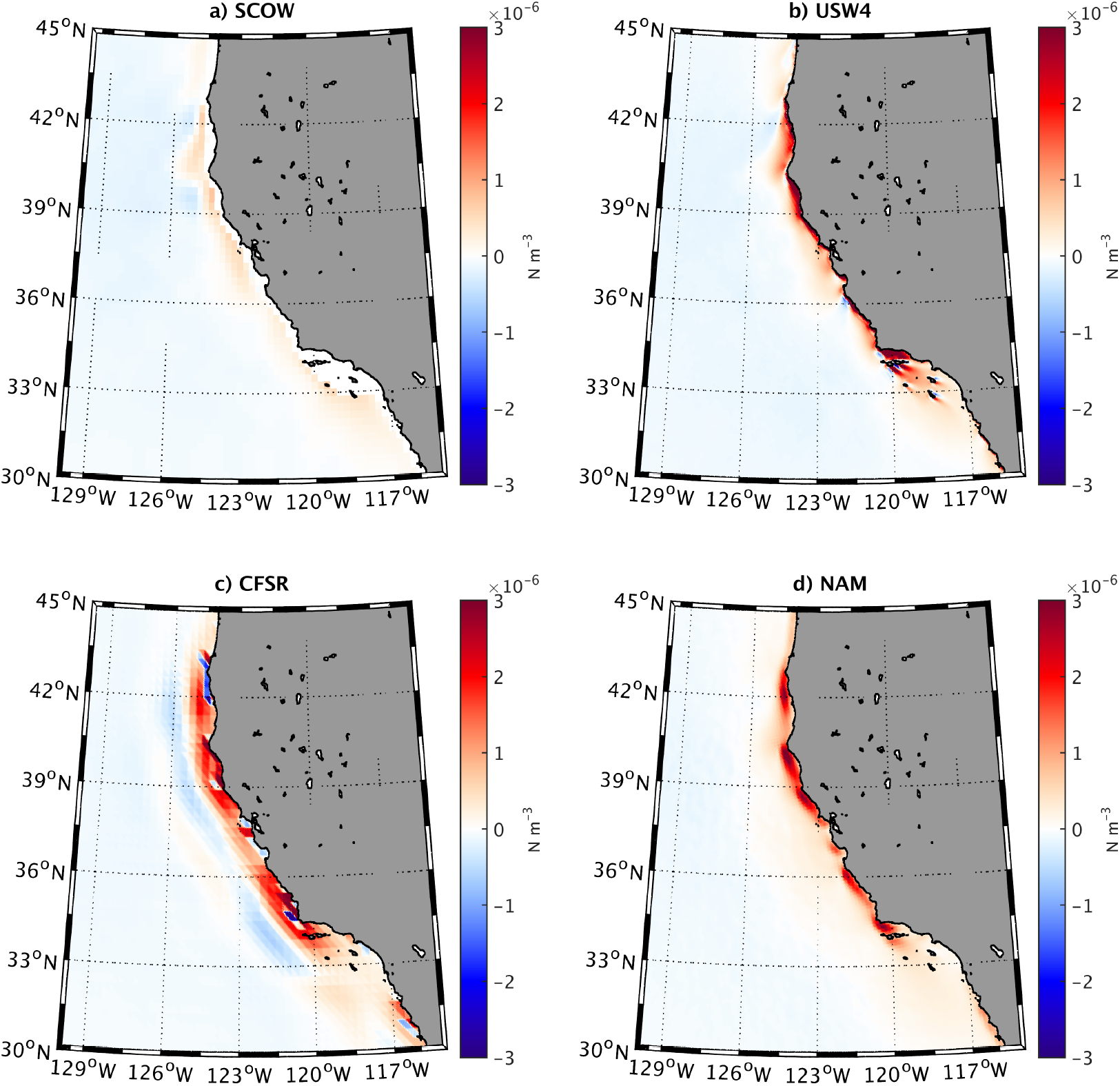
Mean surface stress curl during the upwelling season (Spring and Summer) from (a) QuikSCAT, (b) USW4, (c) CFSR, and (d) NAM. The USW4 atmospheric forcing allows to simulate a wind-drop off, and fine-scale stress curl such as island wind wakes. Note the nearshore blind zone (in white) of QuikSCAT in (a).

Finally, Fig. 6def shows the interannual yearly values of northward surface stress from QuikSCAT and USW4 averaged over the South, Central, and North boxes. Again, the interannual variability is reproduced by the model.

## 4 Oceanic Fields

### 4.1 Surface Layer

To evaluate the performance of the simulated oceanic simulation in terms of seasonal SST, we use the Ostia product as a comparison (Fig. 9), examining both Nearshore and Offshore boxes. In global coupled models, eastern boundary upwelling systems, such as the CCS, are characterized by large SST biases (up to 3°C; *e.g*., Richter (2015)). The origin of these biases is not well understood, but it is likely to be caused by poor representations of the cloud cover, surface wind pattern, oceanic upwelling, and cross-shore eddy heat flux. In USW4 the mean SST large-scale patterns are qualitatively well represented compared to the SST satellite measurements, with warmer waters to the west and south of the domain and colder waters to the north (Fig. 9). The favorable upwelling season (*i.e*., spring and summer) is captured by the simulation, and the upwelling signature is clearly marked in a 30 km wide coastal strip (Fig. 9). The simulated SST has a weak cold bias of 0.5°C over the whole domain in all the seasons, and the coastal water is colder in the model than in the Ostia product by up to 1°C (Figs. 9 and 10). Very nearshore the bias can reach up to 2°, but it is likely due to the limitation of the SST product. Ostia SST has a relatively high spatial resolution of nominally *dx* = 5 km. However, high cloud cover over the upwelling season impedes access to high-resolution data. Therefore, the effective resolution of this product may be similar to that of the microwave satellite products (*dx* = 25 km); this may partially explain the nearshore SST discrepancies between USW4 and the Ostia product. The largest bias is situated in the Southern California Bight where the atmospheric forcing is also less skillful (*i.e*., overestimation of the surface stress and underestimation of the shortwave radiation during summer). Generally, numerical simulations have difficulty representing the southward extension of the cold water south of Pt. Conception (Marchesiello et al., 2003; Capet et al., 2008a; Renault et al., 2016a) (see also Fig. 10). USW4 has at least fair representation of this southward extension; *e.g*., it has a better representation of the SST and of its mean pattern than a climatological solution (or *e.g*., the Veneziani et al. (2009) solution). In particular, in the Central Nearshore box, the climatological solution has a warm bias up to 1.5°C, whereas USW4 has a bias lower than 0.5°C there. Otherwise, the USW4 SST compares well with the measurements, which is likely due to a good representation of the simulated atmospheric forcing (in particular, the cloud cover and wind drop-off) and of the surface currents.

**Figure 9:**
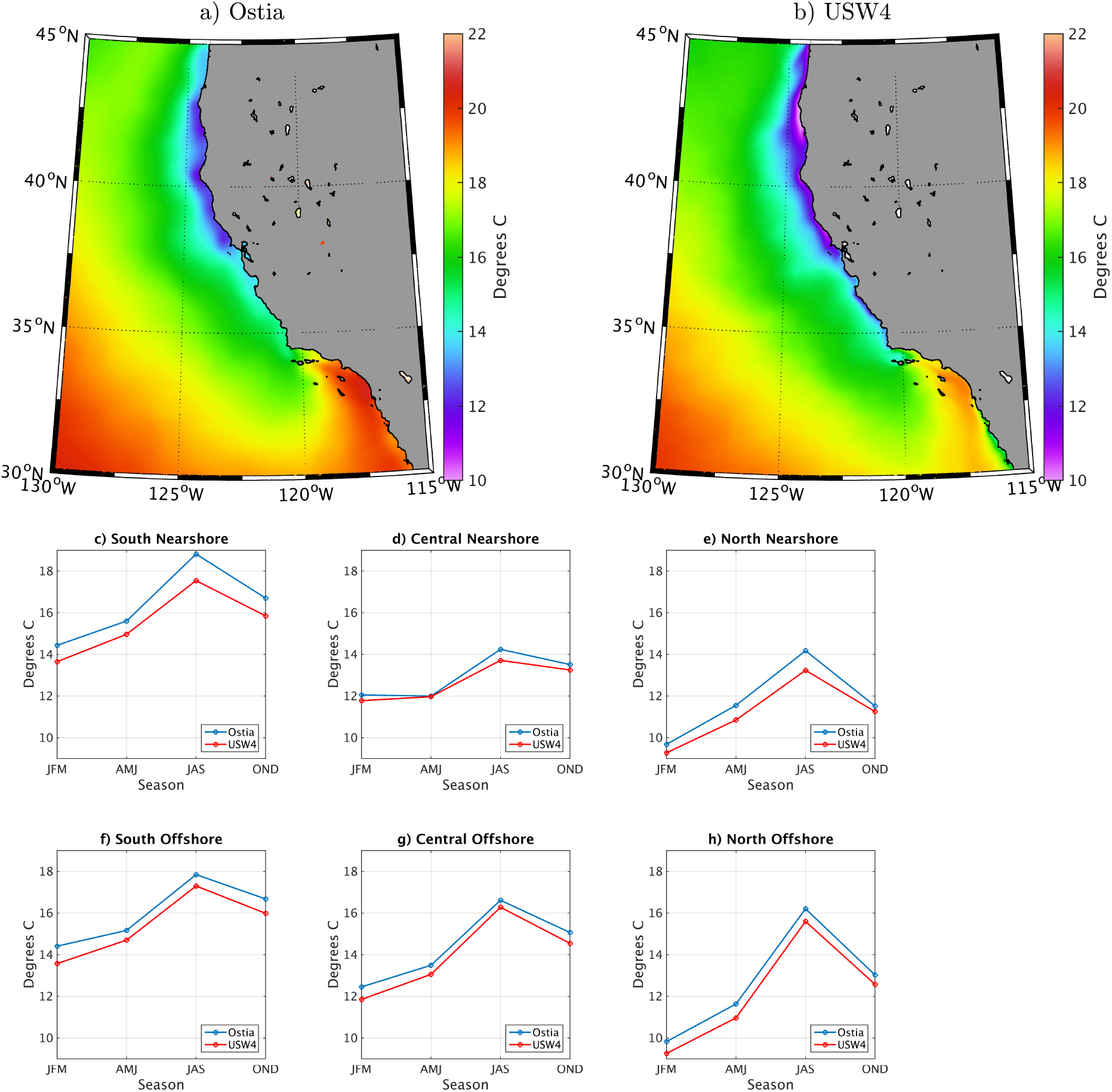
Mean SST [°C] during summer from (a) Ostia and (b) USW4 (1995-2010). Panels (c) to (h) represent the seasonal evolution of the SST over the same period from Ostia (blue) and USW4 (red) and averaged over the Nearshore and Offshore boxes indicated in Fig. 1 or over the whole domain. The SST patterns are well matched in USW4 with a mean negative bias of about 0.5°C.

**Figure 10:**
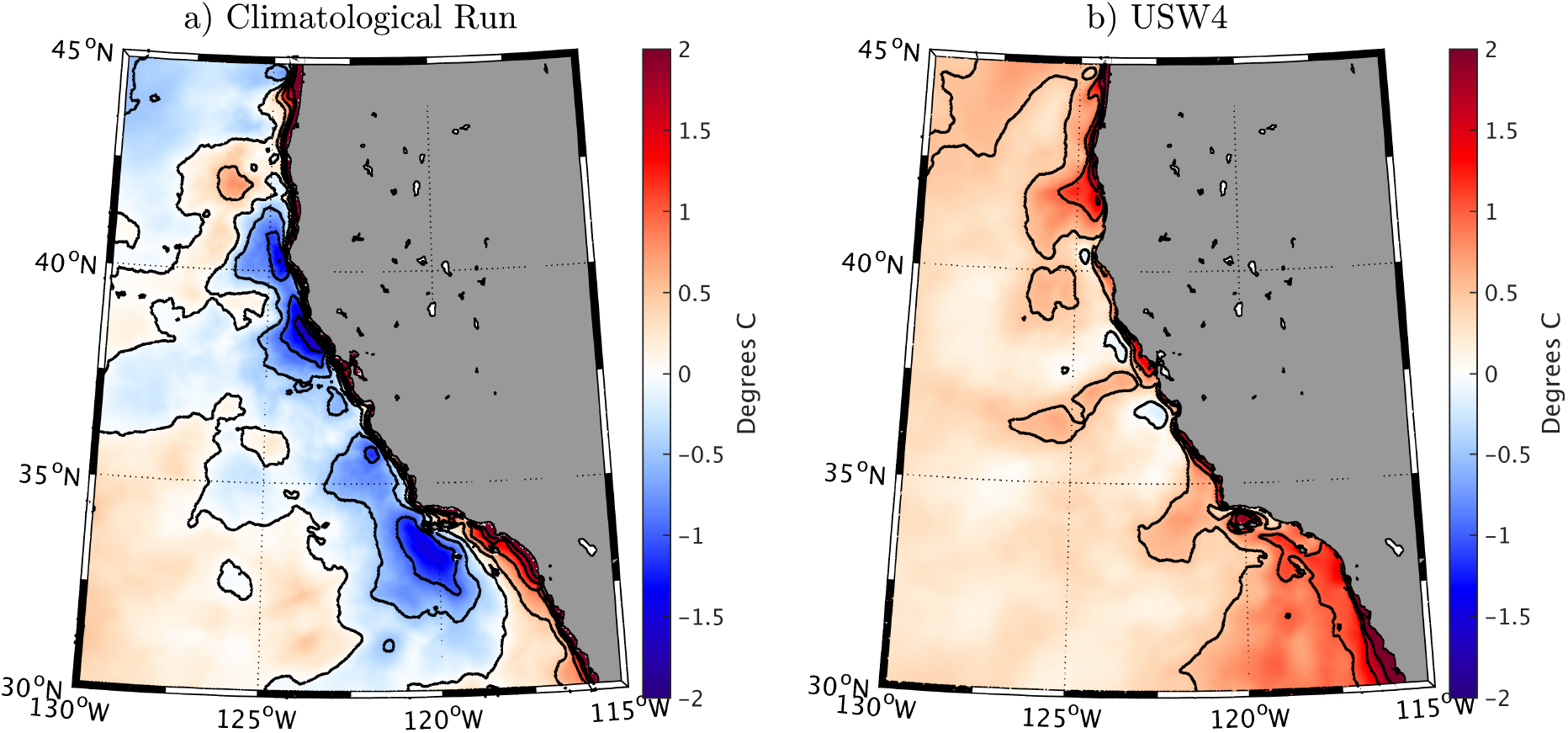
(a) Mean SST differences [°C] during summer between Ostia and (a) a climatological solution (*e.g*., Capet et al. (2008b); Renault et al. (2016a)) and (b) USW4. USW4 has a cold bias (> 0.5 °), in particular over the Southern California Bight; however, it is less biased than the climatological solution (up to 2°C; see text in Sec. 4.1).

Figure 11 depicts the interannual variation of the yearly mean SST in the nearshore region over the South, Central, and North boxes for Ostia and USW4. Consistent with Fig. 3, the interannual variability of the SST is well reproduced by USW4. In particular, the warm anomalies of > 0.5° during 1997 and 2004 and the cold anomaly during 2008 are captured by the model. Similar results are found for the offshore region.

**Figure 11:**
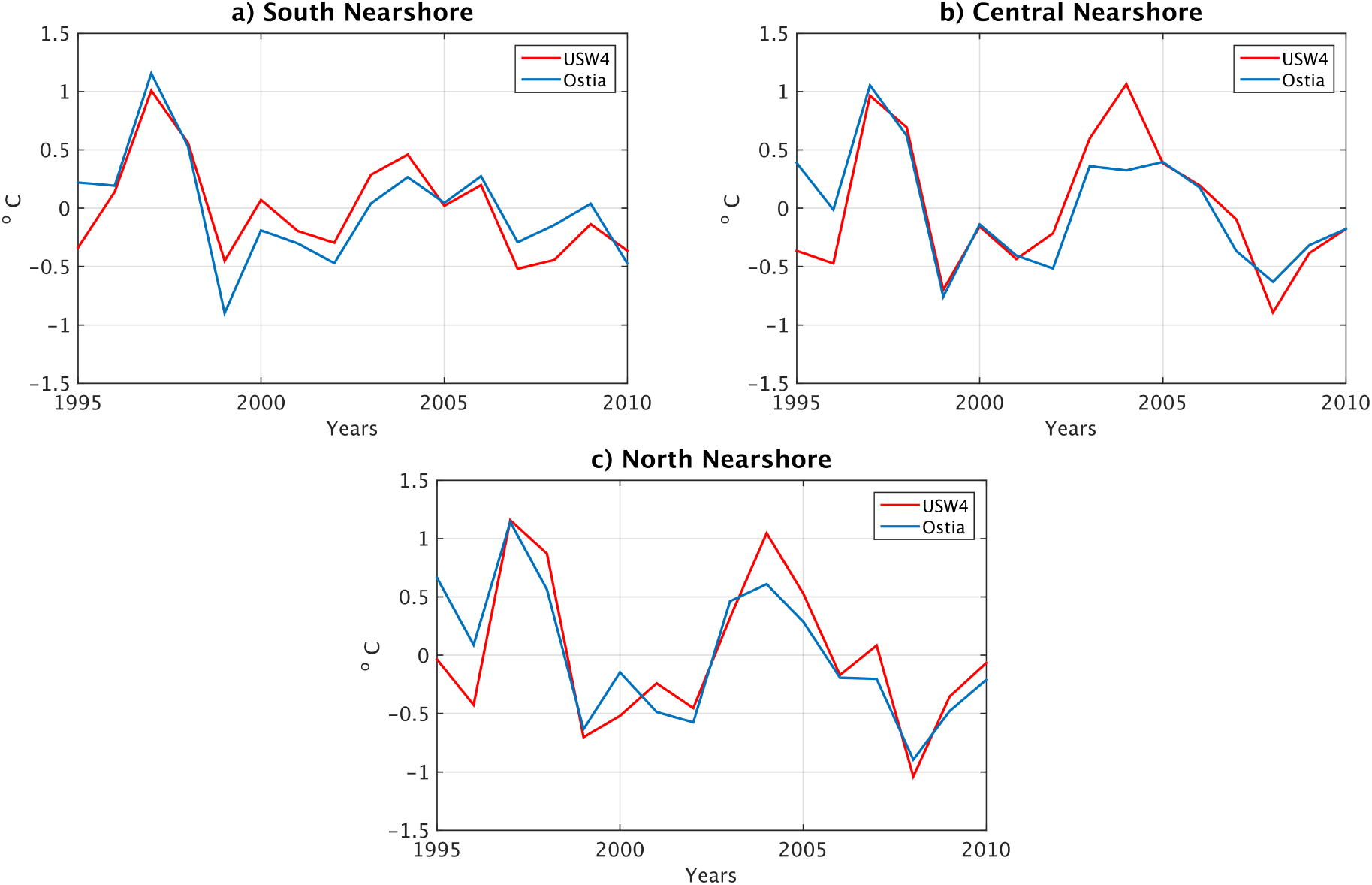
Interannual SST [°C] over the boxes indicated in Fig. 1 from OSTIA and USW4. Similar results are found over the offshore boxes.

Sea Surface Salinity (SSS) is higher offshore in the subtropical gyre with its high evaporation rate and is lower in the subpolar gyre with the higher precipitation. In addition, it decreases weakly near the coast, more so in the north, mainly due to river inflow. We compare the large-scale pattern and seasonal cycle of SSS from USW4 to those from the WOA13 SSS large-scale pattern and seasonal cycle in Fig. 12. Due to a realistic representation of the freshwater flux by USW4 (Sec. 3c), there is good agreement between the simulation and the measurements. However, offshore in central California, the SSS is generally too low with respect to the measurements, with a maximum bias of 0.5 PSU. This is partially explained by the freshwater flux biases in Fig. 5, where the offshore flux is slightly underestimated. However, as discussed in Secs. 2.1 and 4.2, the bias is also partially inherited from the parent solution and its open boundary conditions. Near the Columbia River the USW4 SSS is conspicuously fresher than in WOD13, but the latter is probably horizontally overly smoothed.

**Figure 12:**
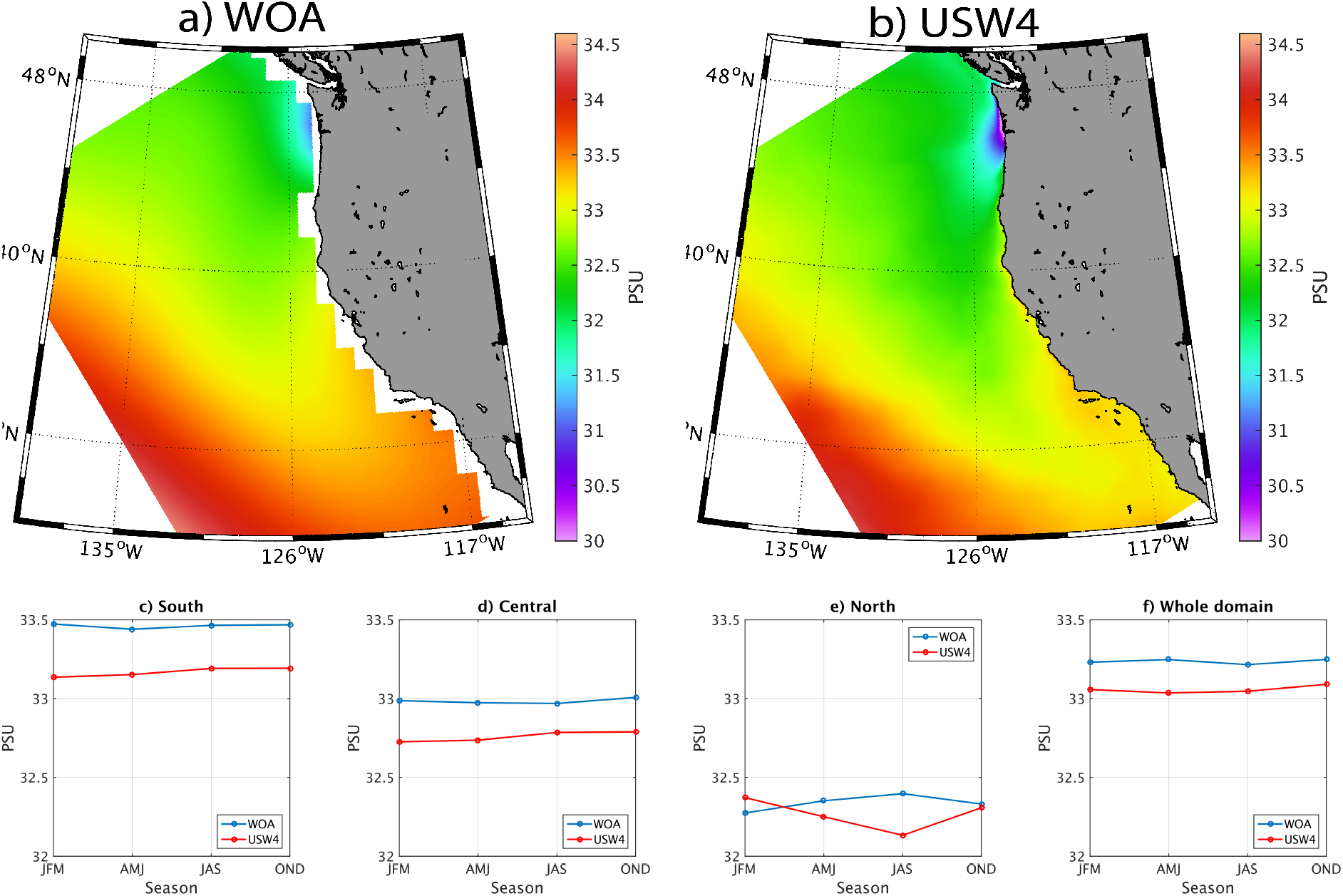
Mean SSS [PSU] from (a) the World Ocean Atlas and (b) USW4 (1995-2010). Due to a realistic freshwater flux, the mean SSS is in USW4 is consistent with the observations despite a small negative bias (up to 0.5 PSU).

In previous studies (*e.g*., de Boyer Montégut et al. (2004)), the Mixed Layer Depth (MLD) definition can be based on different parameters such as temperature, salinity, and density. The MLD is typically defined using a threshold, for which the MLD is the depth at which potential temperature or potential density changes by a specified small value relative to its value near the surface. Here the CARS analysis definition is applied to the daily average temperature and density field in USW4. The MLD is defined using a temperature threshold of ΔΘ = 0.2 ° and Δ*σ_theta_* = 0.3 kg m^−3^. The near-surface reference depth is 10 m. MLD is shallower near the coast due to an uplifted pycnocline, and it is deeper in summer in the offshore subpolar gyre due to its depressed pycnocline compared to the offshore subpolar gyre with its uplifted pycnocline. The CARS MLD and the one estimated from USW4 are compared in Fig. 13. The phase and amplitude of the seasonal cycle are similar in both the model and measurements: a shallowing of the MLD during spring and summer, then a deepening from 20 m to 80 m in winter. The MLD in the model is slightly too deep compared to the climatology; this could be related to the surface forcing and to the KPP parameterization scheme (Large et al., 1994) for vertical mixing of tracers and momentum in ROMS. The vertical mixing that is too deep partially explains the cold bias in the simulated SST.

**Figure 13:**
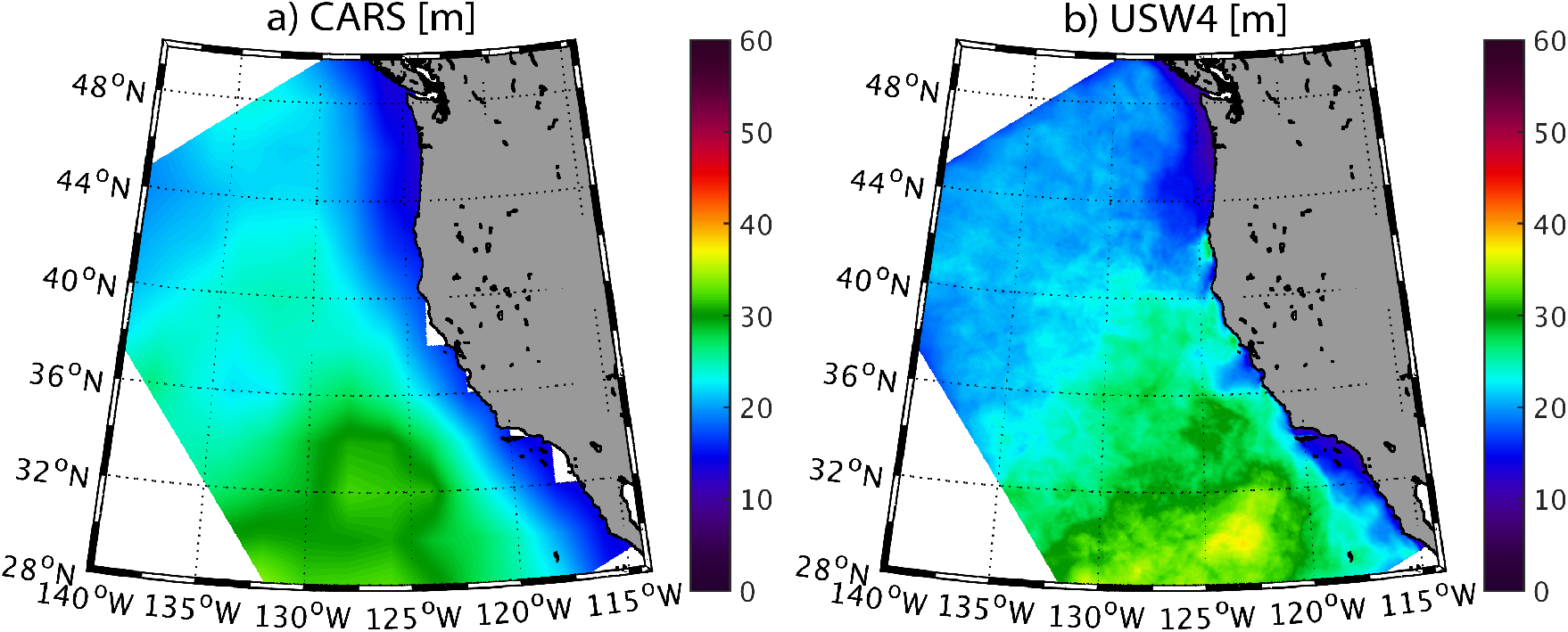
Mean Mixed Layer Depth (MLD) [m] estimated from (a) CARS and (b) USW4 during summer (1995-2010). Panels (c), (d), (e), and (f) represent the seasonal evolution of MLD over the same period from CARS (blue) and USW4 (red), averaged over the boxes indicated in Fig. 1 or over the whole domain. The simulation has a realistic MLD that is deeper farther offshore and during winter, but with a slight overestimation (up to 20 m in the Southern California Bight during winter) that partially explains the cold bias in SST in USW4.

### 4.2 Interior *T* and *S*

The CCS is stably stratified almost everywhere. It has warm temperature and fresh salinity in and above the pycnocline compared to below, and the pycnocline tilts upward toward the coast due to upwelling. Systematic large-scale hydrographic sampling of the CCS was initiated in 1949 by the California Cooperative Oceanic Fisheries Investigations (CalCOFI) program. Along the zonal line 80 of the CalCOFI data (offshore from Pt. Conception, centered on 33°N), the vertical structure of the simulated temperature, salinity, and density are in general agreement with the CalCOFI climatology (Fig. 14). In both the observations and in USW4, isotherms, isohalines, and isopycnals are characterized by a positive cross-shore slope. At the surface, consistent with Fig. 9, the mean SST is well reproduced with biases lower than 0.5 °, which is likely due to a realistic representation of the net surface heat flux. As shown in Fig. 12, the mean salinity in the upper layer is too low with respect to CalCOFI (by 0.2 PSU). Finally, at depth, the mean density field is also realistic; nevertheless, there is a cold temperature bias of 1 ° (a negative density bias) that is partly compensated for by a fresh salinity bias of 0.s PSU (a positive density bias). Similar results are found for the Newport line using the WOD13 dataset (1955-2013, Fig. 14).

**Figure 14:**
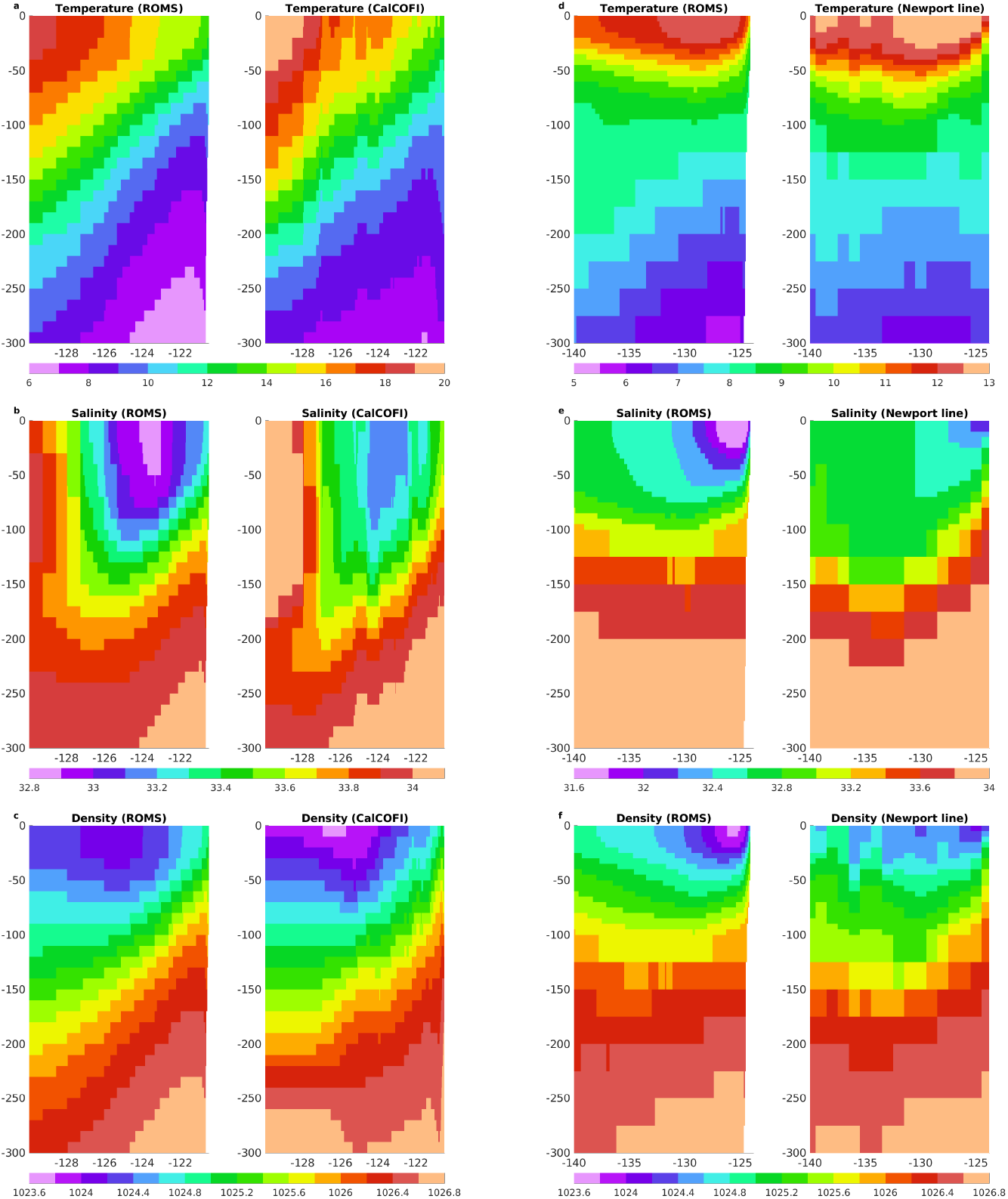
Mean cross-shore section along the CalCOFI lines 80 (left panels, ≈ 33°N) and Newport line from WOD13 (right panels, ≈ 45°N) of (a,d) temperature [°C], (b,e) salinity [PSU], and (c,f) density [kg m^−3^] from USW4 (1995-2010) (left column) and the measurements (period 1955-2013, right column). USW4 has approximately the right cross-shore density slope induced by the wind-driven upwelling. At the surface the salinity is too low with respect to CalCOFI. At depth the density is similar to that in the observations, but there is a cold temperature bias (a positive density bias of ≈ 1°C), partially compensated for by a fresh salinity bias (a negative density bias) of ≈ 0.2 PSU).

Figure 15 shows the mean temperature, salinity, and density biases at 150 m depth. In the first 500 km from the coast, the density is realistic with a very weak bias by less than 0.1 kg m^−3^, and nearshore, where there is more data, the bias is less than 0.05 kg m^−3^. However, consistent with Fig. 14, there is a bias compensation between the temperature and salinity. As suggested by Fig. 14, the temperature and salinity biases are mostly inherited from the open boundary conditions (*i.e*., Mercator fields corrected by WOA) used in the parent-grid solution. Sensitivity tests have been made to reduce these biases. For example, more realistic results are obtained by using Mercator (compared to the Simple Oceanic Data Assimilation (SODA) by Carton and Giese (2008)) as the open boundary condition of the parent simulation and by correcting these data with WOA (not shown). In general, however, the sampling density of measurements in the offshore region is rather small, and we choose not to artificially diminish our large-scale biases by adjusting the boundary conditions within their (considerable) level of uncertainty. Deutsch et al. (2021a) discusses the importance of correctly reproducing the density field in specifying the biogeochemical boundary conditions.

**Figure 15:**
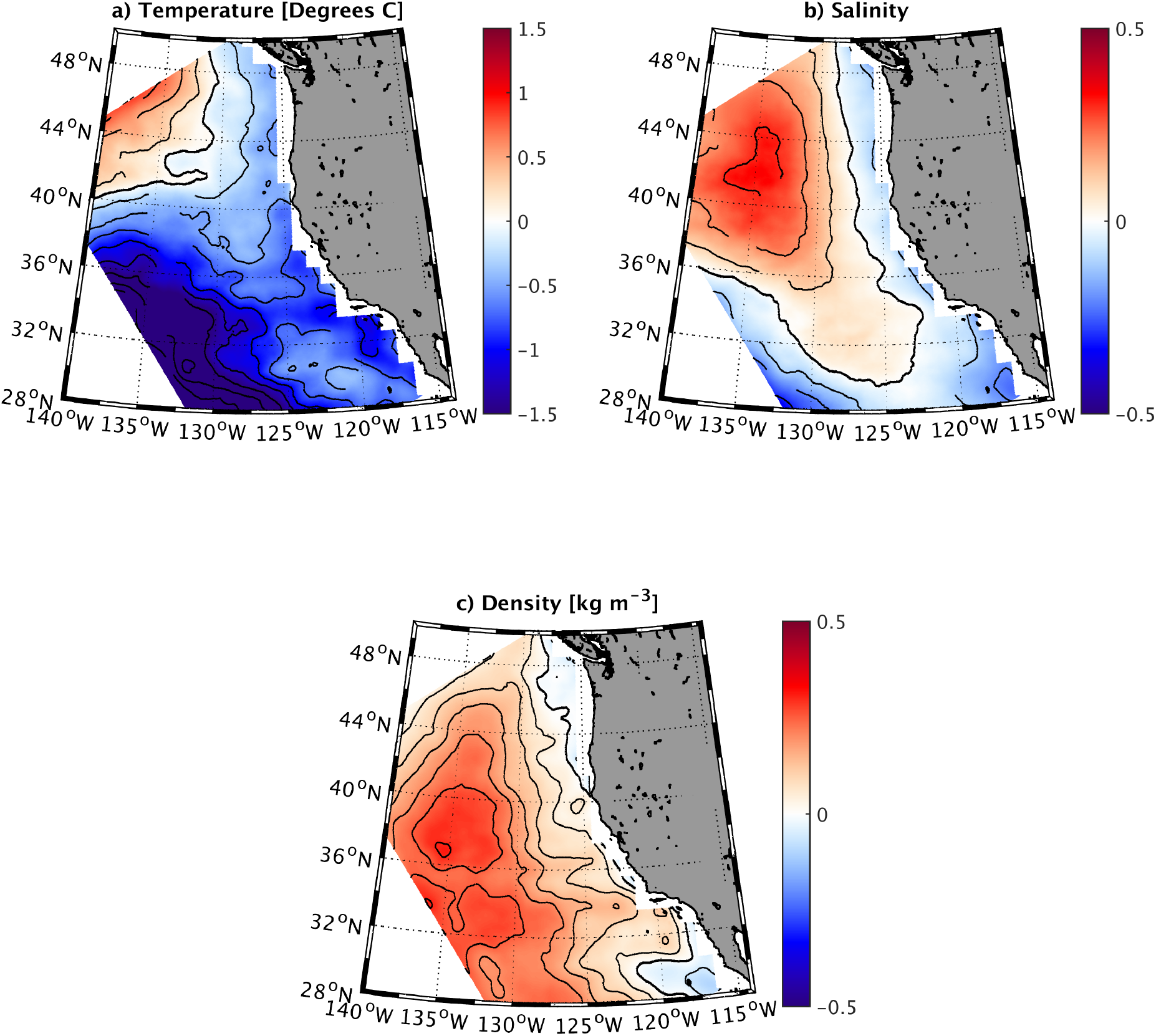
Mean temperature [°C], salinity [PSU], and density [kg m^−3^] differences at 150 m depth between USW4 (1995-2010) and WOA. The contour lines difference isolines, with the thick black line indicating zero difference. In the first 500 km the density at 150 m is realistic, with a very weak bias of less than 0.1 kg m^−3^; nearshore, where there is more data, the bias is less than 0.05 kg m^−3^. However, consistent with Fig. 14, there is a compensation between temperature and salinity biases. Most of the salinity bias enters the domain through the northern open boundary condition.

The temperature and salinity variability in USW4 is furthermore evaluated by comparing their standard deviations (removing the long-term mean) at 150 m to the Word Ocean Database 2013 (WOD13, Fig. 16). In the nearshore region (first 200 km from the coast), as in the measurements, the upwelling has a signature on the water masses: there is a weaker temperature and salinity variability with respect to offshore. Offshore the *T* and *S* variability is mainly due to variations in the pycnocline depth in the subtropical gyres. In USW4 the offshore salinity variability is slightly too weak compared to the measurements; while no doubt part of this discrepancy may be due to model bias (*e.g*., inherited from the parent solution and boundary conditions), it could also partially be explained by uncertainties in the measurements (*e.g*., the salinity variability pattern in WOD13 is somewhat noisy). However, the offshore temperature variability is well reproduced by the model.

**Figure 16:**
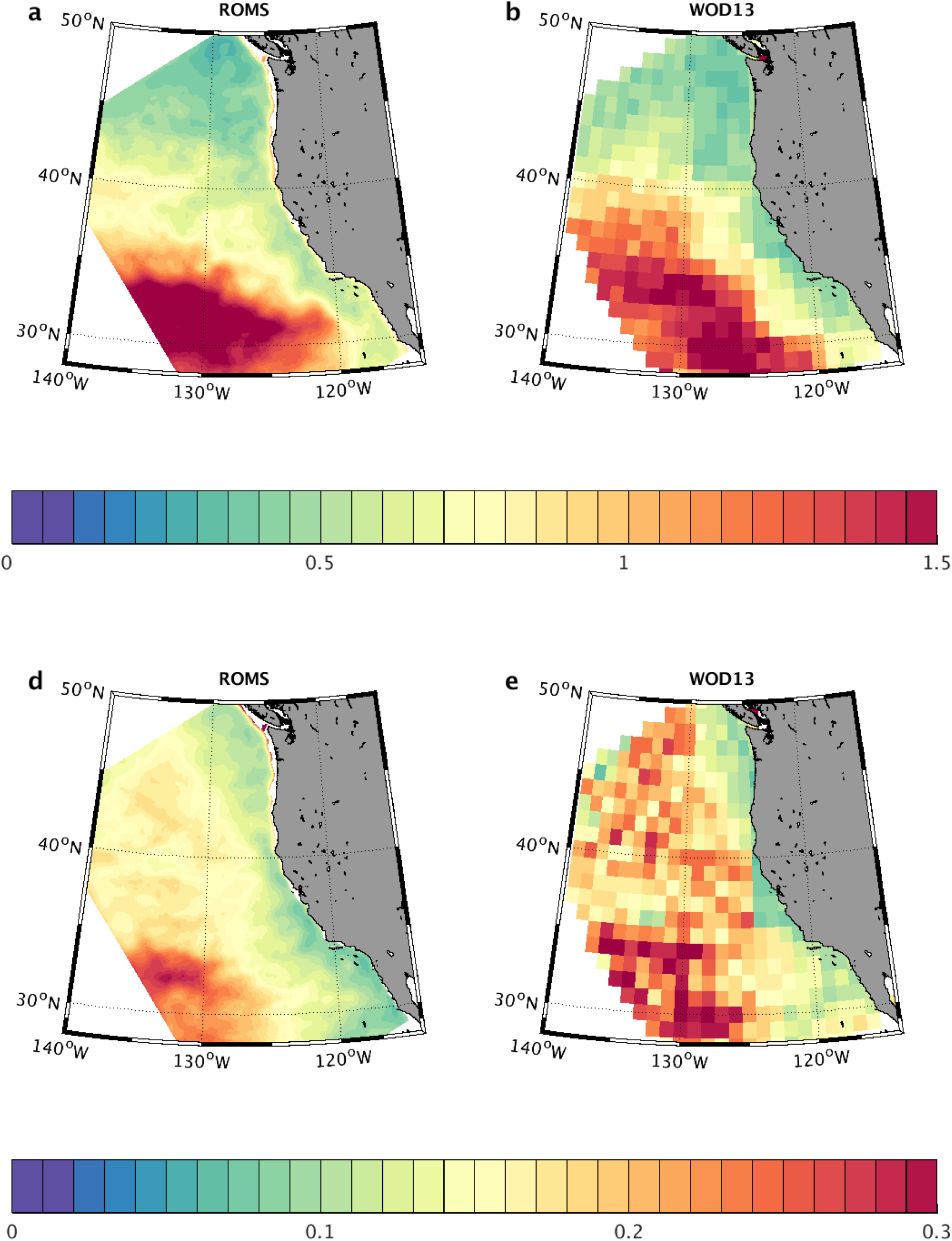
Top panel: Temporal standard deviation of monthly temperature [°C] at 150 m depth for (a) USW4 and (b) (WOD13). Bottom panel: same as the top panel but for the salinity [PSU]. There is a general agreement between the simulated temperature and salinity variability at 150 m depth and the measurements.

Finally, Fig. 17 shows the Potential Temperature - Salinity (TS) diagram as estimated from WOA and from USW4 between 30°-49°N in the upper 1000 m near the coastline (0-100 km offshore). The mean water masses of the CCS are realistic in USW4, although there are cold temperature (≈ 0.5°) and fresh salinity biases (0.2 PSU in the upper layer, maximum bias of 0.5 PSU) consistent with Fig. 14.

**Figure 17:**
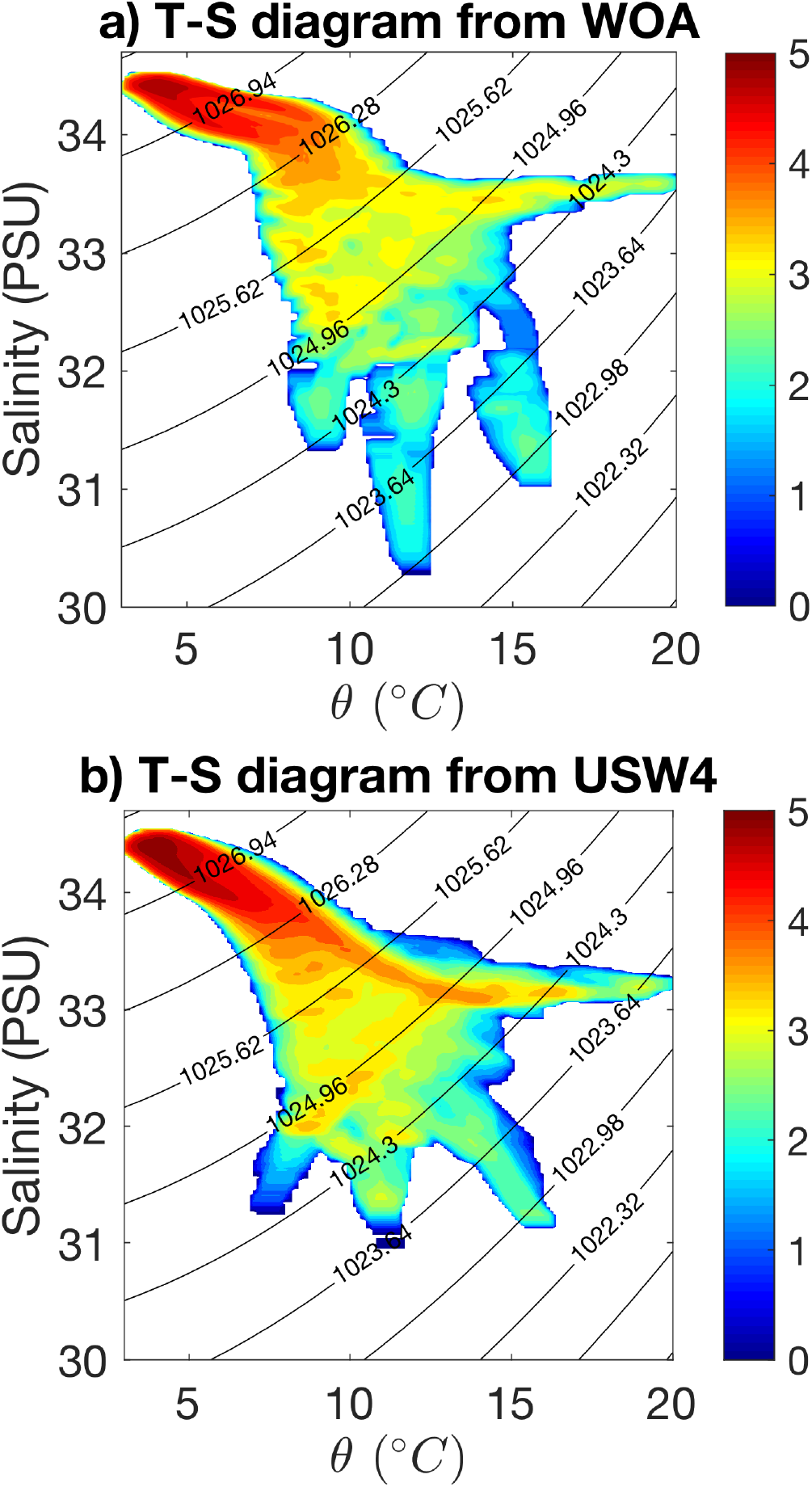
Potential Temperature - Salinity (TS) diagram from (a) the WOA measurements and (b) USW4, for the climatological mean between 30°-49°in the upper 1000 m near the coast (0-100 km offshore). The colorbar shows the number of data points in each (1°C, 0.1PSU) bin on the logarithmic scale, and black contour lines are those of density. The abscissa is potential temperature with the surface as the reference level, and the ordinate is salinity. To obtain the number of data points in each bin, we first obtain the (T,S) dataset in each season, averaged over the years 1995-2010, in the selected region regriding both measurements and USW4 over the a grid with a spatial resolution of *dx* = 4 km in the offshore direction, 0.05°(≈ 5 km) in the along-shore direction, and 20 m in the vertical direction; then, the number of data points in each bin is counted.

### 4.3 Mean Vertical Velocities during the Upwelling Season

Figure 18a represents the mean vertical velocities at 30 m depth (that is near the vertical peak of w nearshore) during the upwelling season as simulated in USW4. Consistent with the literature, the CCS region is characterized by various upwelling cells. The largest vertical velocities reach values greater than 0.5 m s^−1^ on average are located between 42°N and 43°N, 40°N, 39°N, 38°N, 36°N and in the Santa Barbara Channel, *i.e*., near capes, complex orography, and coastline that strengthen the wind. The associated subseasonal variability is shown in Fig. 18b. It reveals a large variability reaching up to 2 m s^−1^ in the nearshore region and a non-negligible variability offshore of 0.5 ms^−1^. Such a variability is associated with wind bursts that induce intense upwelling and larger turbulent heat fluxes (Renault et al., 2009) but also to the mesoscale activity. The interannual variability is also relatively large (Figure 18c) with values greater than 0.5 m s^−1^ nearshore, and that is mainly associated with the interannual variability of the wind (see e.g., Fig. 6).

**Figure 18:**
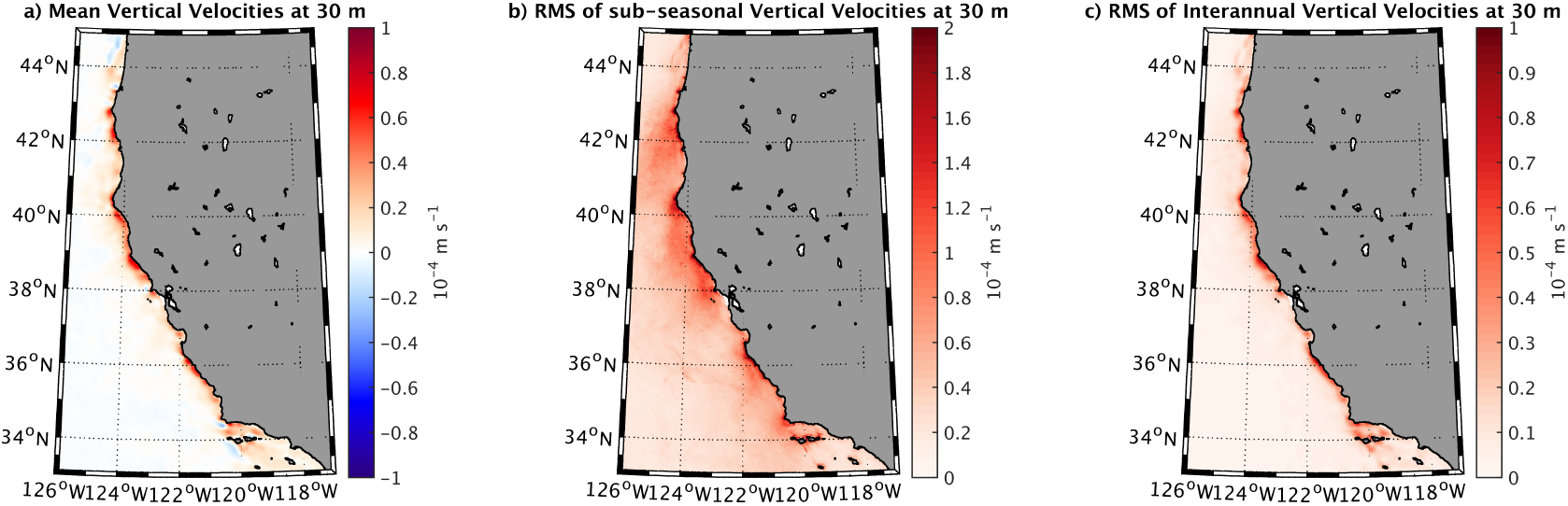
Vertical velocities [m s ^−1^] during the upwelling season at 30 m depth as simulated by USW4. (a) Long-term mean, (b) subseasonal variability, (c) interannual variability.

### 4.4 Mean Sea Surface Height and Current

The SSH (Sec. 2c) from the 16 years of USW4 is shown in Fig. 19ab, along with measurements from the 1/4°resolution CNES-CLS13 dataset (Rio et al. (2014), Sec. 2d). The spatial distribution and amplitude of the simulated SSH is in good agreement with the measurements. The mean Sea Surface Height (SSH) in the CCS is depressed at the coast due to the southward geostrophic current, and it further decreases poleward due to the equatorward wind stress. The main differences between the model and measurements are located along the coast. Such discrepancies can be attributed partially to the Nearshore box width (50 km) which is unresolved in the satellite data (Ducet et al., 2000; Rio et al., 2014). The negative cross-shore SSH slope is reproduced by USW4. Interestingly, the alongshore standing eddies are much less evident in USW4 and in the CNES-CLS13 dataset than in drifter measurements (Centurioni et al., 2008) or the model by Marchesiello et al. (2003), likely because of the longer time averaging used here. To better highlight the presence of standing eddies, an Empirical Orthogonal Function (EOF) analysis is applied to the 16 years of daily SSH from USW4 after removing the mean state (Sea Level Anomaly, SLA) over a nearshore region shown in Fig. 19cd. The obtained modes have therefore to be interpreted as the variation of the circulation with respect to the mean state. Here focus is mainly done on the modes that are characterized by the presence of standing eddies. The first EOF mode (not shown) explains 34.8% of the variance and depicts the steric contribution. The second EOF mode (not shown) explains 22.3% of the variance, it represents the seasonal variation of the surface currents (*e.g*., southward intensification during the upwelling season, see below). More interestingly, the third and fourth modes explain 12.2% and 6.5% of the variance, respectively. Figure 19cd depicts their spatial patterns and Figure 19ef their associated temporal variations and spectrum. Both Mode 3 and 4 reveal the presence of standing eddies from 200 km offshore in particular around 39°N and 36°N. Spectral analysis of the associated series reveals significant energy peaks at ≈ 630 and 10 day^−1^. The 95is estimated by a Markov red noise (Gilman et al., 1963). The 10 day^−1^ peak likely represents wind burst that induce a modulation of the currents. Mode 3 also reveals a variation of the nearshore current that is likely responsible of the frequency peak near 25 and 90 days^−1^.

**Figure 19:**
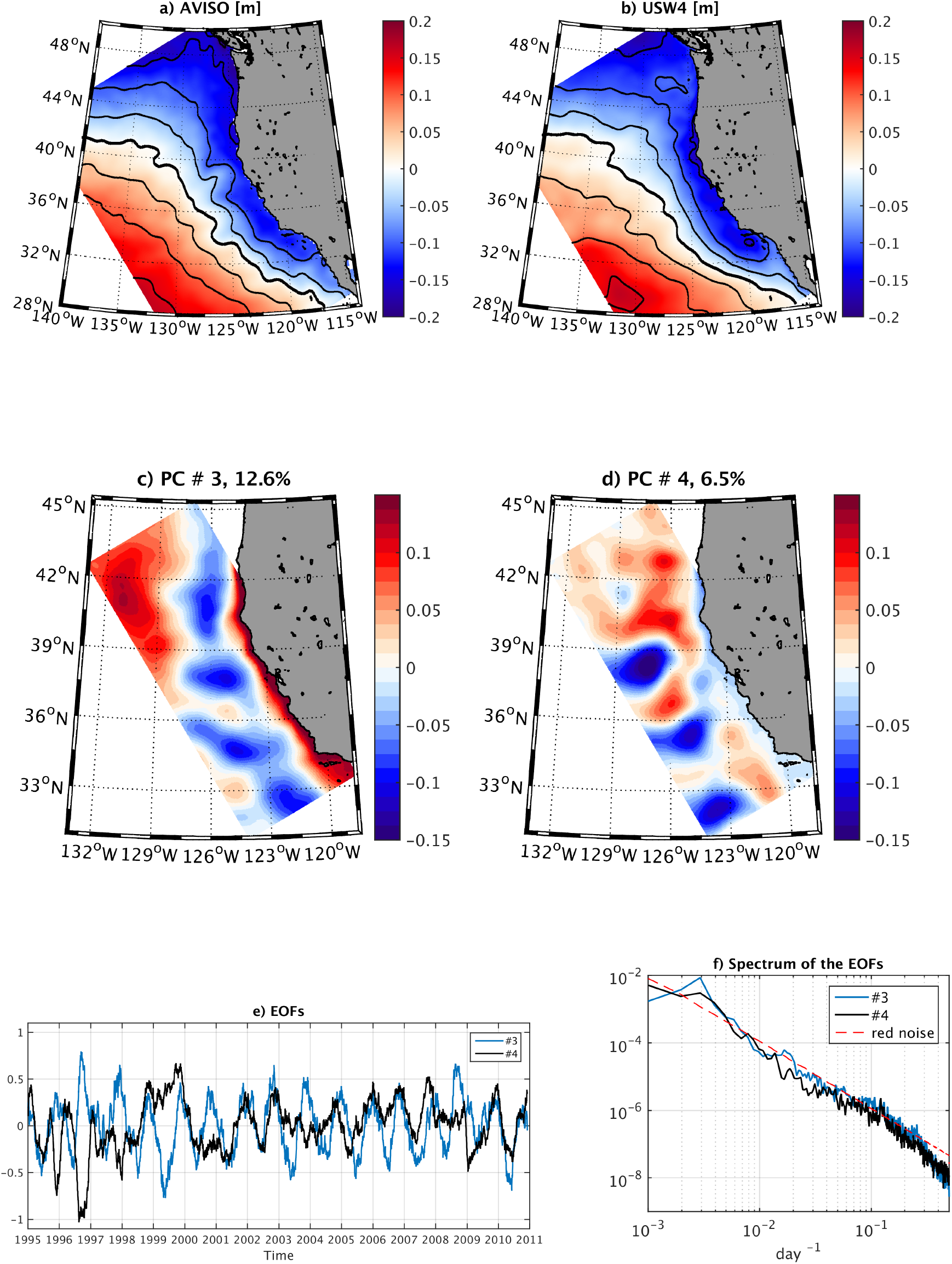
Mean Sea Surface Height (SSH) [m] from (a) AVISO and (b) USW4 over the period 1995-2010. Contours show 0.05 m increments of SSH, and the thick black line represents the local zero reference height contour. The simulation reproduces the mean SSH and its offshore gradient. c-d-e) represent the third and fourth mode of the Empirical Orthogonal Function (EOF) decomposition of the Sea Level Anomaly and the associated timeseries. f) Spectrum of the EOF timeseries. Standing eddies can be identified on the EOF pattern modes.

The CCS exhibits broad-band variability. On time and space scales larger than the mesoscale, most of the variability is partly extrinsic to the region, reflecting the larger-scale seasonal, interannual, and decadal climate signals that have regional manifestations along the U. S. West Coast (Chhak and Di Lorenzo, 2007; Di Lorenzo et al., 2009; Chenillat et al., 2012; Meinvielle and Johnson, 2013; Davis and Di Lorenzo, 2015). To give a sense of this variability, Fig. 20 shows the whole-coast history of the sea-surface height anomaly (SLA) and depth of a pycnocline isopycnal over the hindcast period (See also Fig. 22 in the biogeochemical companion (Deutsch et al., 2021a)). The along-coast coherence of SLA is striking, as is the regularity of its primarily seasonal oscillation. Its amplitude increases to the north because the amplitude of the seasonal cycle of the alongshore wind and current increase as well. A rapid poleward propagation speed ≈ 2.5 m s^−1^ is apparent on average, as has been extensively analyzed previously (Chelton, 1984; Spillane et al., 1987). The interannual variability amplitude is a modest fraction of the seasonal one, most of the time, but the 1997-98 ENSO event is particularly prominent (Kosro, 2002; Ryan and Noble, 2002; Lynn and Bograd, 2002).

**Figure 20:**
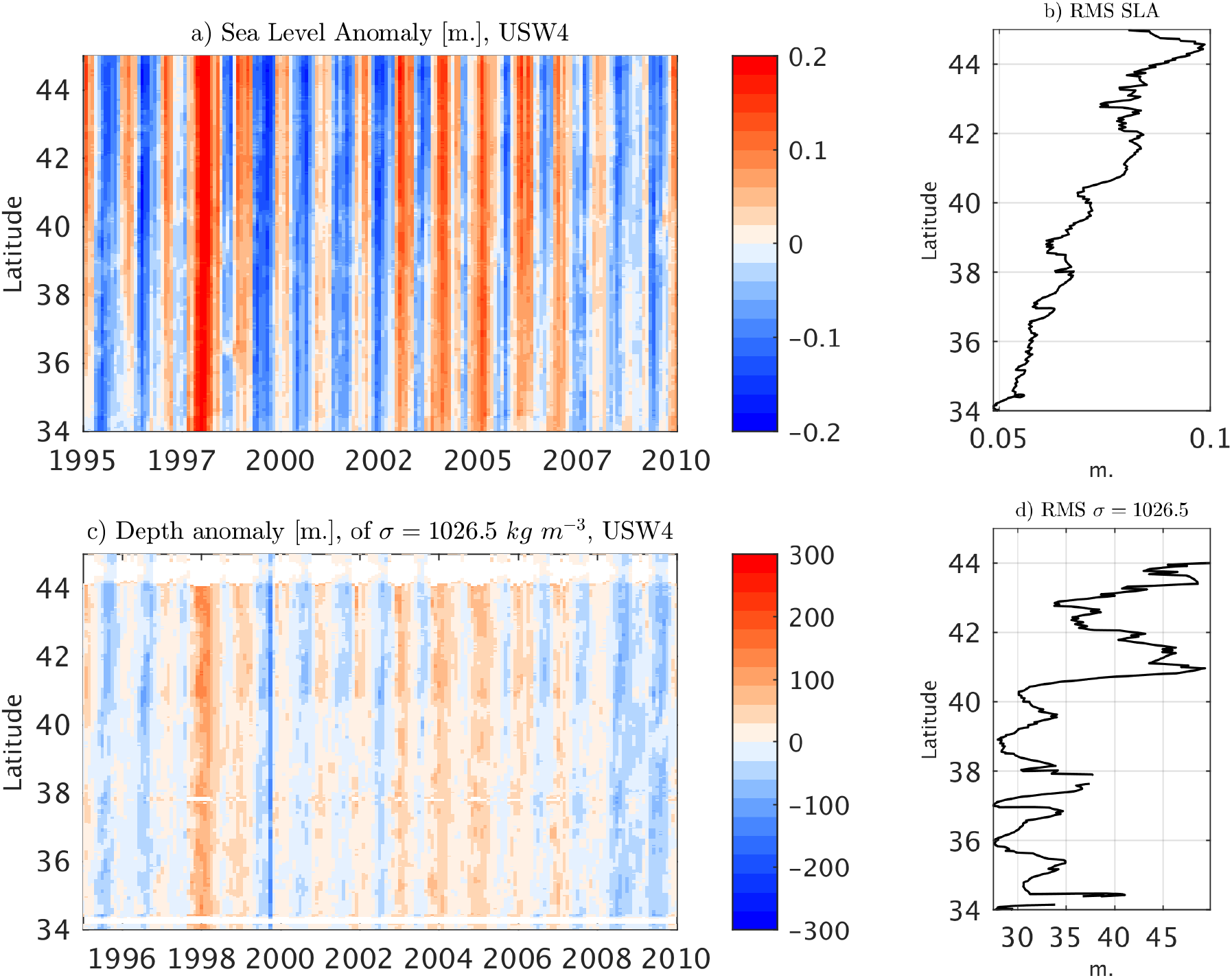
Hovmöller diagrams (latitude and time) for (a) SLA [m] and (c) depth of the *σ_θ_* = 25.6 isopycnal surface [m] at a distance ≈ 50 km offshore. On the right of each plot is the corresponding RMS latitude profile with respect to the temporal variability.

The pattern in pycnocline depth is somewhat more complex, although the temporal correlation with SLA is evident (*C* ≈ 0.8, more or less uniformly along the coast). The amplitudes of both quantities have an increasing trend to the north, but the depth anomaly exhibits more modulation in amplitude than the SLA with moderate drops at particular latitudes that are related to interruptions in the path of the California Undercurrent (CUC) along the coast (Chen et al., 2021).

The spatial pattern of the geostrophic current estimated from the observed and simulated SSH is also in good agreement (not shown). Figure 21a shows the nearshore meridional surface current during the upwelling season. The nearshore surface current is broad and generally equatorward, as observed (Swenson and Niiler, 1996). It reaches values up to 0.2 m s^−1^. The nearshore surface current properties exhibit strong latitudinal variability (values from ≈ 0 to 0.2 m s^1^). The maximum amplitude of the current is situated along the coast and near capes, *i.e*., where the wind is more intense. Figure 21bc represents the subseasonal and the interannual variability of the meridional surface current. The subseasonal variability of the surface meridional current during the upwelling season is large, reaching values up to 0.3 m s^−1^, i.e., larger than the seasonal averages. Such a variability is mainly associated with the mesoscale activity and with wind bursts that modulate the Ekman transport and the upwelling-associated geostrophic currents.

**Figure 21:**
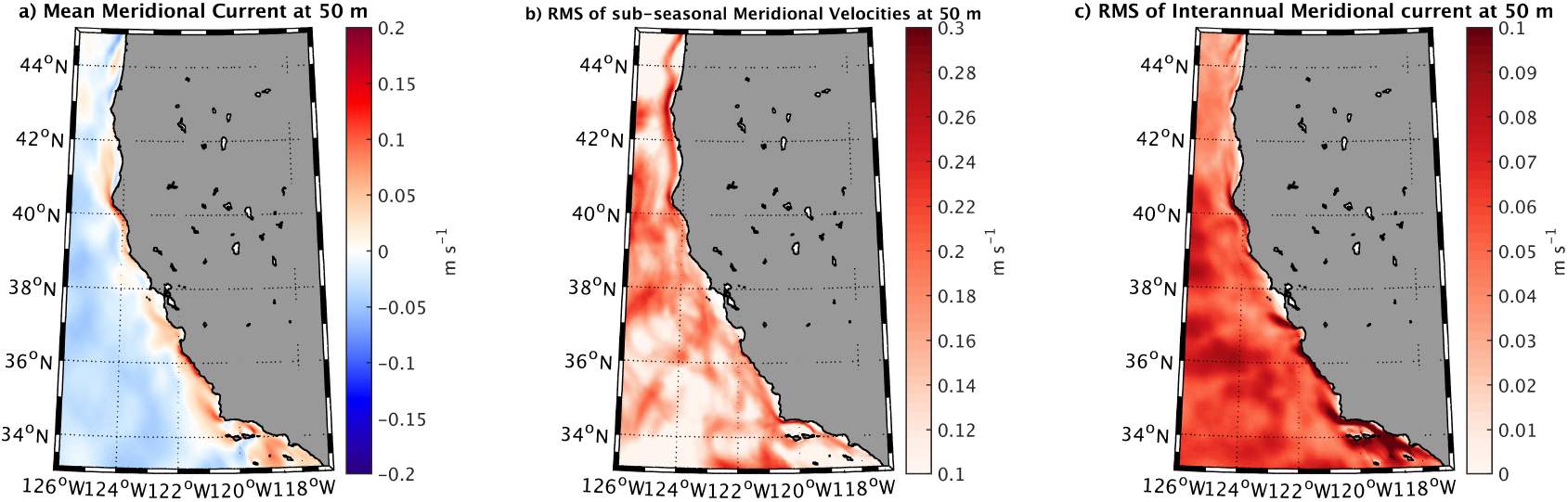
Meridional current at 50 m depth [m s^−1^] from USW4. (a) Long-term mean during the hindcast period, (b) subseasonal variability, (c) interannual variability. Note that the surface nearshore current is southward.

Figure 22 depicts coastal cross-shore sections of the seasonal meridional current averaged along CalCOFI glider line 66.7 (Rudnick et al. (2017) and https://spraydata.ucsd.edu/climCUGN/), *i.e*., near San Francisco Bay, for winter, spring, summer, and fall. As reported in the measurements, the coastal CCS during the upwelling season is characterized by an equatorward surface current with a mean velocity of 0.1 m s^−1^ overlying the CUC. The CUC, one of the major components of the CCS, is a poleward flow in the upper hundreds of meters near the U. S. West Coast. It transports warm and salty equatorial equator poleward and plays a significant role in the local heat, salt and biogeochemical budgets. Quite a few studies exist characterizing the features and exploring the dynamics of the CUC (*e.g*., McCreary et al. (1987); Lynn and Simpson (1987); Pierce et al. (2000); Gay and Chereskin (2009); Molemaker et al. (2015); Rudnick et al. (2017)). Using USW4, Chen et al. (2021) assess the CUC dynamics and show that topographic form stress is a significant northward acceleration effect for this current both in its mean and low-frequency variability.

**Figure 22:**
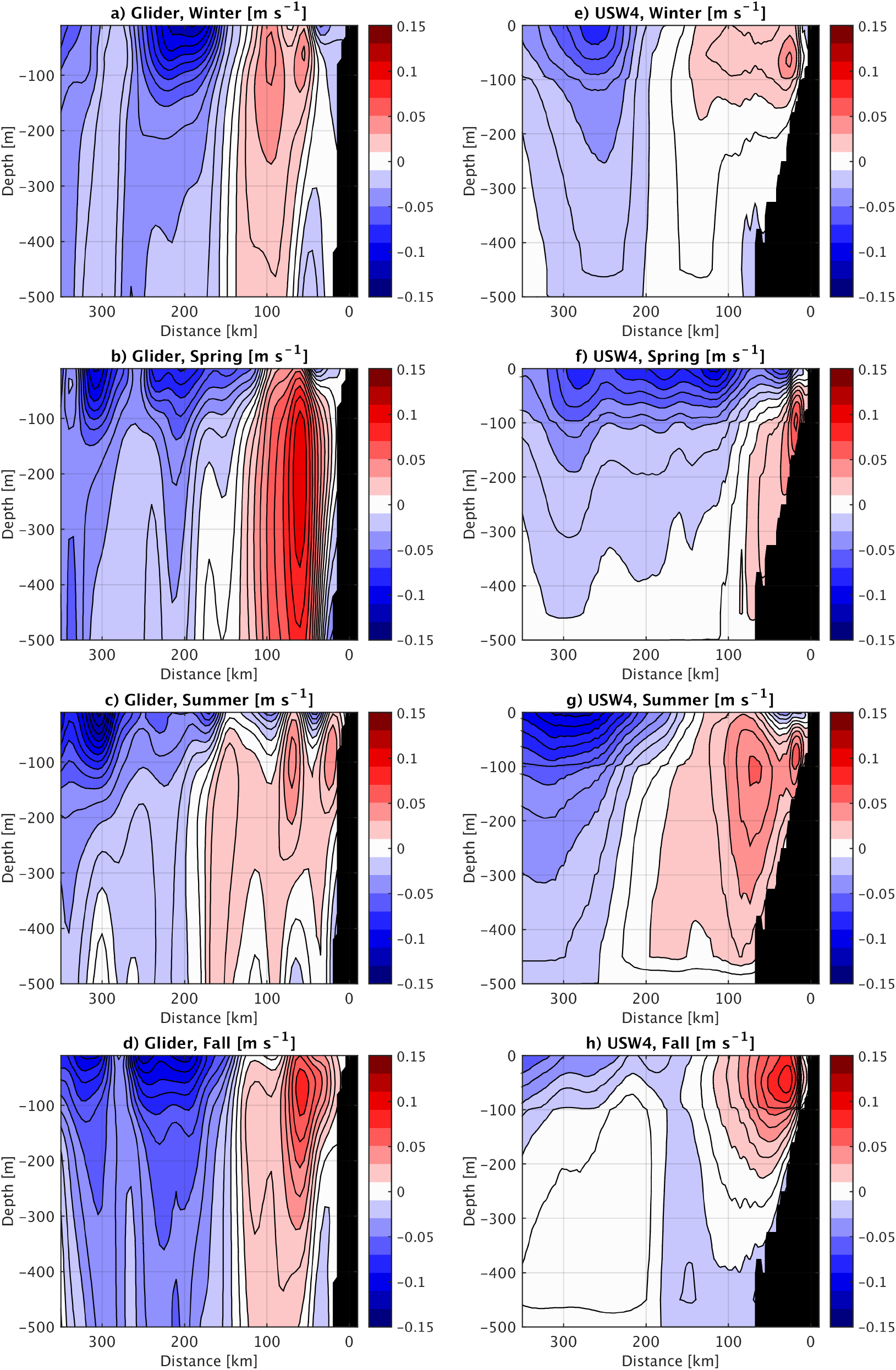
Seasonal means of the meridional geostrophic current [m s^−1^] along line 66.7 estimated from the gliders (abcd) and USW4 (e,f,g,h). Note the definition of the seasons differs from that in Rudnick et al. (2017).

The CUC structure is well represented in USW4 for the winter, summer, and spring seasons. In both USW4 and the observations, the core of the CUC is relatively shallow during winter and fall (50 m depth) and is deeper during summer (100 m depth). In summer, a surface equatoward current (0.05 *ms*^−1^) overlies the CUC whereas in fall, the CUC outcrops the surface, reversing poleward the surface current. In Winter, the CUC still outcrops the surface but the surface current remains equatoward. These is also an indirect validation of the simulated wind drop-off, as a poor representation of the slackening of the wind toward the coast (as in CFSR) may cause occasional surfacing of the California Undercurrent (CUC) (Renault et al., 2016a) through Sverdrup dynamics: a positive surface stress curl produces a barotropic poleward flow that adds to the coastal baroclinic flow (McCreary and Chao, 1985; Lynn and Simpson, 1990; Marchesiello et al., 2003). The spring season is characterized by a bias in the representation of the CUC characteristics. In the observations, the CUC has intense velocities (up 0.1 *ms*^−1^) and has a core reaching a depth of 200 m (in particular in June, not shown). In USW4, the CUC remains weak (velocities of 0.05 *ms*^−1^) and its core it not deep enough (100 m depth). Note that the definition of the seasons here differ from that of, Rudnick et al. (2017). This affects the interpretation of the seasonal cycle and of this bias as the main discrepancy occurs in June (summer in Rudnick et al. (2017), spring in this study).

To further assess the realism of the USW4 CUC, we characterize the interannual and the subseasonal variabilities of the meridional geostrophic currents along line 66.7 from the gliders and from USW4 Both gliders and USW4 reveal an interannual variability associated with current anomalies reaching up to 0.05 m s^−1^ (not shown). The subseasonal variability induced anomalies larger than 0.15 m s^−1^ (not shown). It is mainly associated with displacement of the CUC related to remote forcing and local wind forcing and with the presence of eddies.

## 5 Higher Frequency Variability

### 5.1 Mesoscale Activity

The geostrophic surface EKE is estimated over the period 1995-2010 from AVISO (Ducet et al., 2000) and from USW4 (Fig. 23ab). For the model the EKE is computed using low-pass filtered geostrophic velocities with a Gaussian spatial filter with a 36-km half-width and a temporal smoothing of 7 days as an approximation to AVISO’s resolution (Chelton and Schlax, 2003) (More precise comparisons could be made using simulation fields processed in the same way as altimetric measurements.). The relevant EKE conversion rates (Sec. 2c) in USW4 are also evaluated over the period 1995-2010 (Fig. 23c-e). Consistent with Strub and James (2000); Marchesiello et al. (2003) and Renault et al. (2016d), the mean CCS circulation is unstable and generates mesoscale eddies primarily by baroclinic instability (Strub and James, 2000; Marchesiello et al., 2003) while the *K_m_K_e_* conversion is a secondary term. In both observations and USW4 The simulated EK has its largest values a couple of hundred km offshore and exhibits a wide decay zone further offshore (Fig. 23). This pattern is due to the combined influences of Ekman transport, eddy dispersion, and the eddy killing effect of the current feedback, with an overall similarity to AVISO and the literature (*e.g*., Capet et al. (2008a)). The overall EKE amplitude is about right in USW4, but there are some biases in the USW4 spatial pattern, mainly that the EKE is too large in the near-coastal region, and the EKE is too large in the Southern California Bight, which seems most likely due to errors in the atmospheric forcing. Part of the discrepancy may be due to model bias, *e.g*., the current feedback induces a dampening of the mesoscale activity, but the wind response induces a partial re-energization. In USW4, following Renault et al. (2016d), the *s_w_* coefficient is used to mimic the wind response to the current feedback. Figure 23f depicts the temporal evolution of the surface EKE domain average from USW4 and from the coupled and uncoupled simulations used in Renault et al. (2016d). USW4 has comparable level of energy as the coupled simulation that includes the wind response to current feedback (EXP3), indicating the parameterization used partly allows to re-energize the mesoscale currents. *s_w_* is taken as spatially and temporally constant, which could induce, for example, a re-energization that is too strong in the nearshore region.

**Figure 23:**
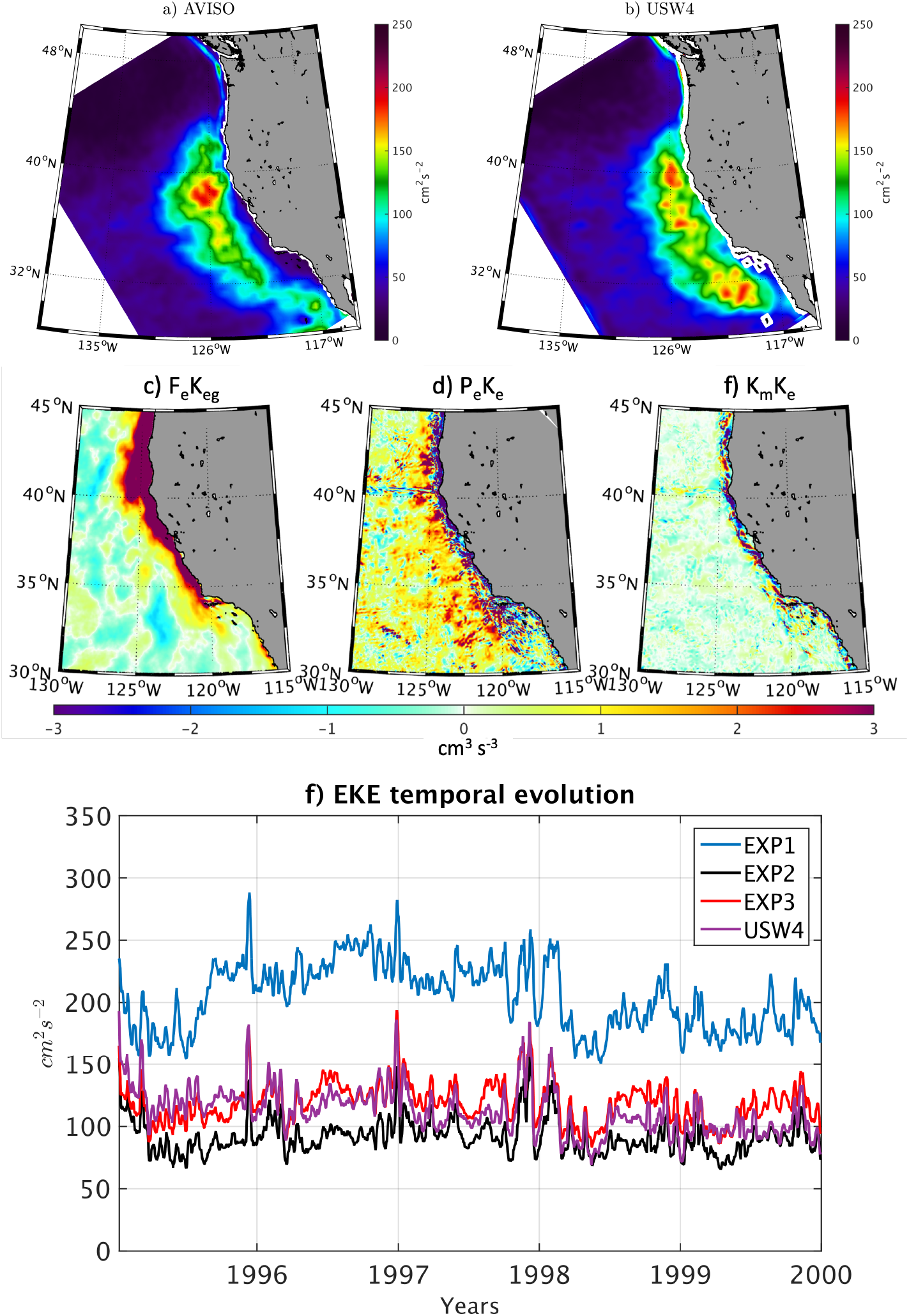
Mean geostrophic surface EKE [cm s^−2^] estimated from (a) AVISO and (b) USW4. The EKE of USW4 is computed using low-pass filtered geostrophic velocities (a Gaussian spatial filter with 36-km-half-width) as an approximate match to AVISO’s spatial resolution. (c)-(e) Geostrophic eddy wind work (*F_e_K_eg_*), baroclinic conversion (*P_e_K_e_*), and barotropic conversion from the mean flow (*K_m_K_e_*) [cm^3^ s^−3^] in USW4 over the period 1995-2010. (f) Temporal evolution of the surface total EKE averaged over the whole domain from USW4 and the simulations from Renault et al. (2016d): EXP1 is a coupled simulation without current feedback, EXP2 is a forced simulation that uses the relative wind to the oceanic motions but without a parameterization of the wind response, EXP3 is a coupled simulation with current feedback.

Figure 24a depicts a cross-section of the EKE averaged between 35°N and 40°N. It reveals that the EKE is large from 200 m depth to the surface and from the coast to ≈ 800 km offshore. The EKE is characterized by a peak of 200 cm^2^ *s*^−2^ at 200 km from the coast and at the surface and slowly decays in the offshore direction while rapidly decreasing at depth with values of less than 75 cm^2^ *s*^−2^. *P_e_K_e_* associated vertical structure is shown in Fig. 23b. It reveals that most of the positive values of *P_e_K_e_* occur in the first 50 m depth, from the coast to 100 km offshore. Finally, a mean cross-shore profile between 30°N and 45°N is estimated for *P_e_K_e_*, *K_m_K_e_*, and *F_e_K_eg_* (Fig. 24c). The geostrophic eddy wind work *F_e_K_eg_* profile is also estimated using a QuikSCAT product (Bentamy and Fillon, 2012) and AVISO, but only over the available QuikSCAT period (2000-2009) (Due to the QuikSCAT and AVISO coastal accuracy issues, the *F_e_K_eg_* value over the first 50 km off the coast is not shown.). Consistent with the measurements, *F_e_K_eg_* is positive in the nearshore region and then becomes negative offshore, deflecting energy from the oceanic geostrophic eddy currents to the atmosphere and thus dampening the offshore eddies. USW4 deflects slightly more energy offshore than the measurements (by 10% averaged over the offshore area). This could be due to estimation errors in the measurements, but it might also be due to biases in the atmospheric and oceanic simulations, with an overestimation of the EKE reservoir (more energy to be deflected) and a biased estimation of *s_w_* when estimating the surface stress. For example, *s_w_* has a seasonal cycle (Renault et al., 2017, 2020) and depends on the atmospheric parameterization of the marine boundary layer (Renault et al., 2016d), and these dependencies are not included in these hindcast simulations. This may explain the overestimation of the offshore EKE and the overestimation of the eddy life in EXP3 shown by Renault et al. (2016d).

**Figure 24:**
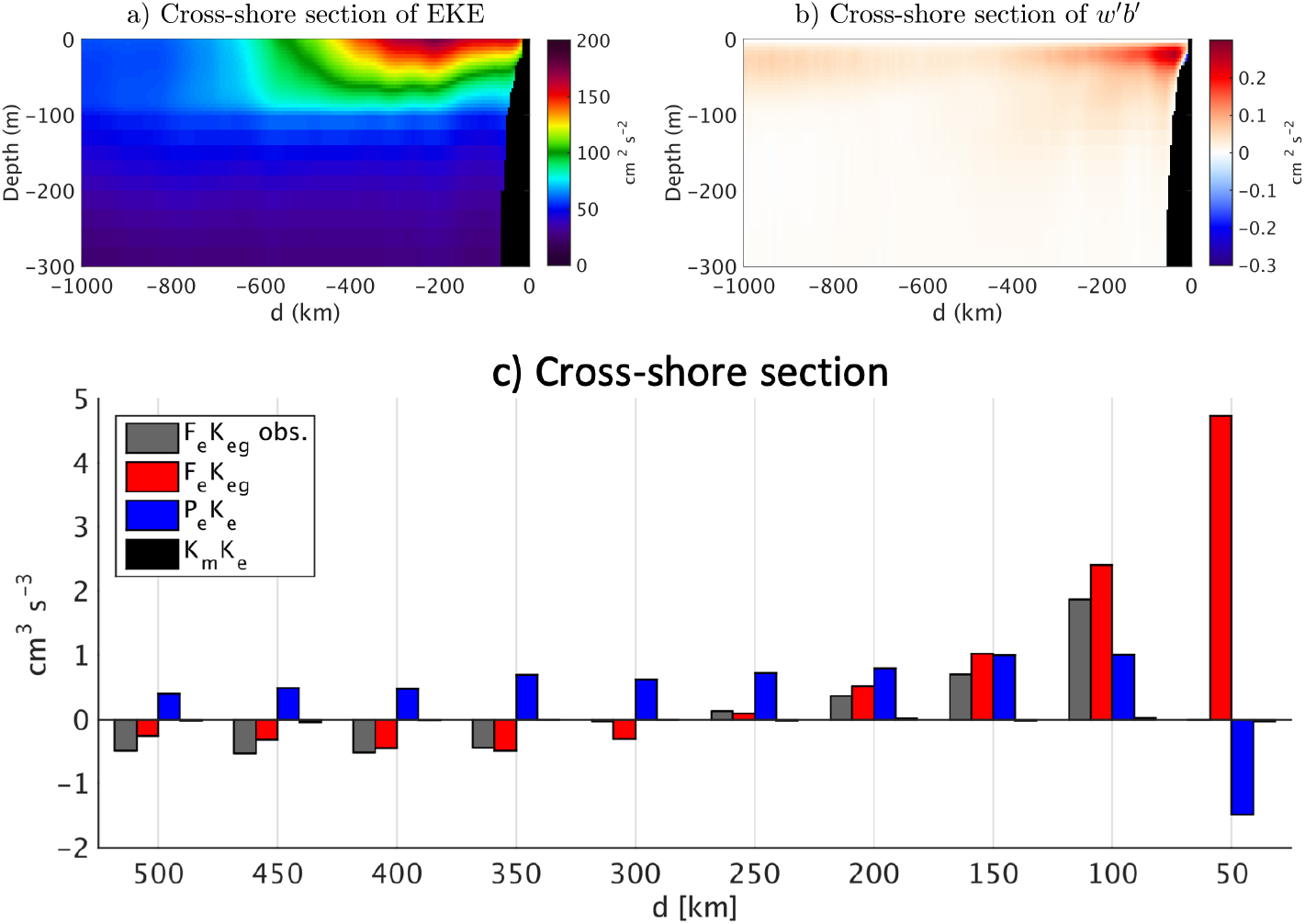
(a)-(b) Cross-shore sections of the mean EKE and *P_e_K_e_* in USW4 averaged between 35°N 40°N. (c) Cross-shore bins of *F_e_K_eg_*, *P_e_K_e_*, and *K_m_K_e_* averaged over 50 km intervals between 30°N and 45°N. The geostrophic eddy wind work (F_e_*K_eg_*) is estimated from USW4 (red) and from measurements (QuikSCAT and AVISO; gray). For the first 50 km off the coast the observational estimate is not shown because of coastal contamination. The baroclinic conversion is the main energy generation term. The eddy wind work *F_e_ K_eg_* is positive nearshore and deflects energy from the ocean to the atmosphere offshore while dampening the mesoscale activity

Figure 25a shows the seasonal cycle of the EKE as estimated from AVISO and from the USW4 low-pass filtered geostrophic velocities with a Gaussian spatial filter with 36-km half-width (solid line) and 28-km-half-width (dashed line). Consistent with *e.g*., Amores et al. (2018), the EKE estimated from AVISO (and from USW4 filtered data is likely to be underestimated by a factor of 2. However, USW4 realistically simulates the seasonal evolution of the EKE, with larger EKE values in summer and fall. The EKE seasonal cycle is mainly driven the seasonal variability of *P_e_K_e_* (not shown). Finally, Fig. 23f reveals a large interannual variability of the EKE, especially in the South and Central boxes.

**Figure 25:**
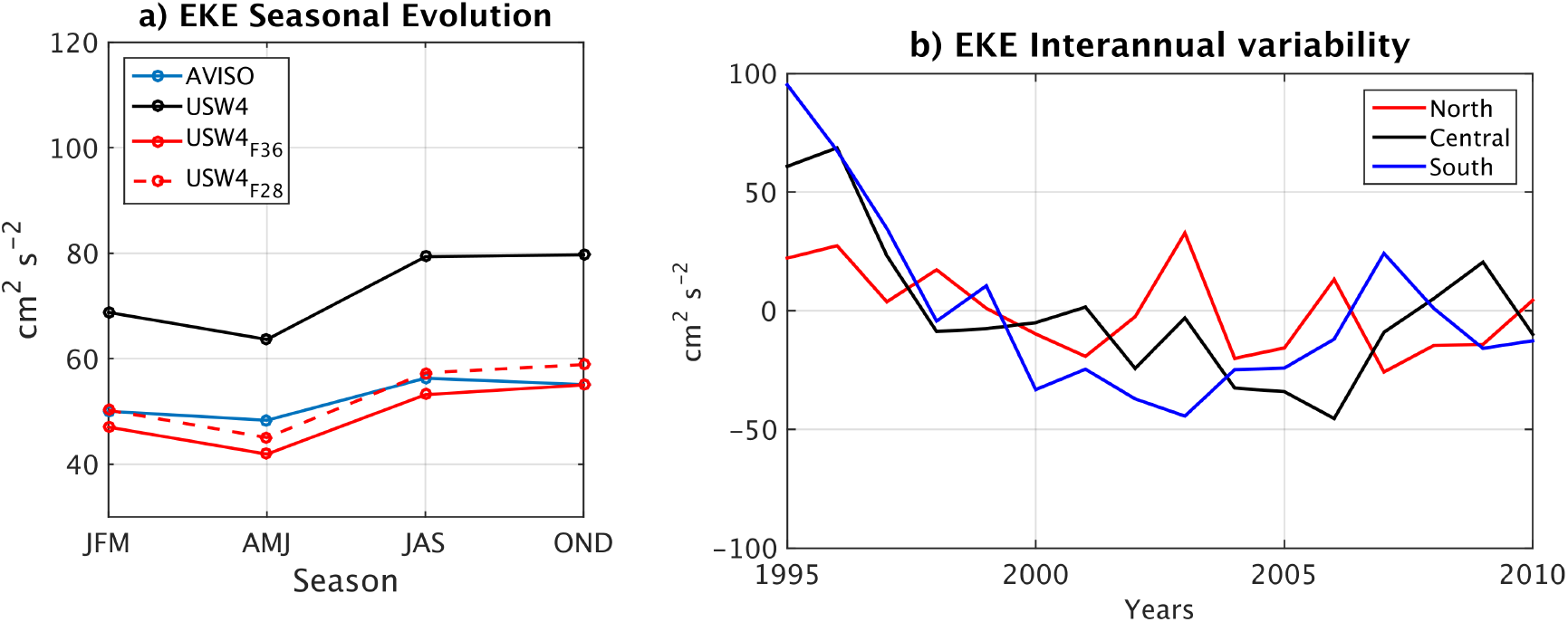
(a) Seasonal evolution of the mean EKE averaged over the whole domain as estimated from the measurements and from the USW4 original filtered geostrophic velocity (based on two different filter half-widths: 36 km (solid line) and 28 km (dashed line)). (b) Interannual variability of the EKE averaged over the boxes indicated on Fig. 1. The current feedback to the atmosphere dampens the eddies and thus allows the simulation to have a realistic EKE level, albeit with not quite the same spatial pattern around the Southern California Bight. Pointwise sampling errors are up to ≈ 5 cm^2^ s^−2^, estimated using a bootstrap method (Efron and Tibshirani, 1985): the mean EKE is computed 100,000 times using random samples from the distribution, and the uncertainty is then defined as ± the standard deviation of these values.

### 5.2 High-Frequency Wind Forcing

In this section the oceanic impact of the synoptic wind variability is assessed. In a bulk formula the surface stress has a quadratic dependence on the wind. As a result, time-varying winds contribute to the time-mean surface stress. It is well known that neglecting high-frequency winds can induce large errors in the surface stress estimate (Esbensen and Kushnir, 1981; Gulev, 1994; Wu et al., 2016). Recent studies show those errors can cause large biases in kinetic energy transfer between the atmosphere and ocean (Zhai et al., 2012; Zhai, 2013). In particular, Zhai et al. (2012) using a global oceanic model showed the mean wind work increases by 70% when using a 6-hourly wind update instead of a monthly one. A few previous studies assess the role of high-frequency atmospheric wind in determining the oceanic circulation. They find that it can lead to an increase by about 50% in both the mean wind work and the EKE, as well as a strengthening of the wind-driven subtropical gyre by about 10-15% (Holdsworth and Myers, 2015; Wu et al., 2016; Condron and Renfrew, 2013).

To determine the importance of high-frequency wind forcing in the CCS, three additional experiments have been carried out for the period 1995-1999 using a 6-hourly-averaged wind forcing (6H), and a daily-averaged wind forcing (1D). The mean alongshore surface stress and the EKE averaged along a coastal band 100 km wide are illustrated in Fig. 26 and summarized in Table 1. From a 1-hourly (*i.e*., USW4) to 6-hourly wind forcing, the surface stress is only slightly impacted by 3%. When estimating the stress using 1D, it is slightly underestimated by 6%. This is, for example, the error made by the QuikSCAT daily products for the CCS. From USW4 to 6H and 1D, the surface current is slightly reduced by 7% and 13%, respectively.

**Figure 26:**
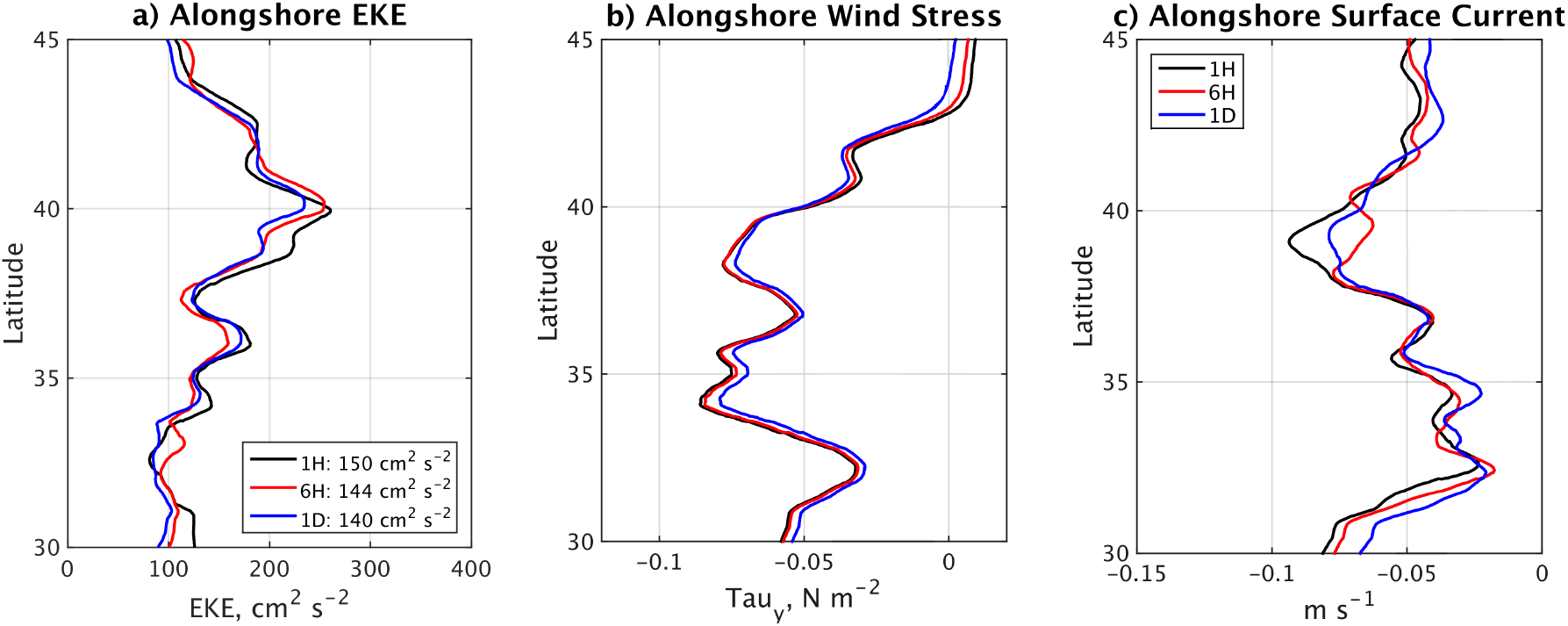
Influence of high-frequency wind forcing on the oceanic surface currents and the surface stress. (a) Annual mean alongshore EKE estimated over a coastal band of 100 km width in USW4 with 1 hr (1H), 6 hr (6H), and 1 day (1D) wind update intervals over the period 1995-1999. (b) Same as (a) but for the mean alongshore wind stress. (c) Same as (a) but for the mean alongshore surface current.

**Table 1:** Mean EKE, surface stress, and alongshore surface current averaged along a coastal band 100 km wide for USW4 (1H forcing) and the three additional experiments using a 6-hourly-averaged wind forcing (6H), and a daily-averaged wind forcing (1D).

| | $EKE_{along} [\text{cm}^2 \text{s}^{-2}]$ | $\tau_{along} [\text{N m}^{-1}]$ | $V_{along} [\text{cm s}^{-1}]$ |
| --- | --- | --- | --- |
| 1H | 150 | -0.049 | -5.3 |
| 6H | 144 | -0.049 | -4.9 |
| 1D | 140 | -0.047 | -4.6 |

The error in wind work has important consequences on the mesoscale activity. As shown in Fig. 23, the main sources of EKE are the baroclinic energy conversion and the eddy wind work. By reducing the mean input of energy *F_m_K_mg_* and the shear of the alongshore current, the absence of the high frequency component of the wind leads to a reduction of the baroclinic conversion rate, *P_e_K_e_*, by 1%, and 4% from USW4 when the wind is temporally smoothed to 6H and 1D, respectively. The positive coastal *F_e_K_eg_* is also reduced by 3%, 8%, and 70% (not shown). As a result the mean alongshore EKE is reduced by 4%, 7%, and 62% in wind-smoothings of 6H and 1D compared to USW4 (Table 1). A simulation forced by a daily atmospheric forcing likely under-estimates the EKE by 7%. An oceanic model could also be forced by a monthly stress (estimated from averaging the hourly stress and thus accumulating the nonlinear effect of synoptic wind); such a simulation would neglect the negative wind work *F_e_ K_eg_*, leading to an overestimation of the EKE by 60% (Fig. 23).

Along a coastline, the cross-shore Ekman transport is proportional to the surface stress, *T_E_* = *τ_alongshore_/ρf*. Thus, the underestimation of the stress by neglecting the high frequency wind leads to a similar underestimation of the transport. The mean Ekman transport is reduced by 2%, and 6% from USW4 to more smoothly varying winds with 6H and 1D averages, respectively.

A striking difference between USW4 and the other simulations is the level of activity in the inertial currents (Fig. 27). By neglecting the hourly stress the inertial currents are much weaker with 1D wind forcing. This is confirmed by the spectrum of the alongshore current: USW4 has a large peak of energy around 18 hours that is not reproduced by 1D. This is consistent with *e.g*., Zhai (2017), who found that almost all the energy flux from the wind to near-inertial motions in the mid-latitude North Pacific and Atlantic are due to a mesoscale atmospheric system with scales less than 1000 km; a high frequency forcing is deemed to be required to represent them. Finally, the lack of inertial currents in 6H and 1D can be seen through the eddy ageostrophic wind work (F_e_*K_ea_*) estimated over the 5 years of simulation from USW4, 6H, and 1D. The *F_e_K_ea_* work is underestimated by 20% by neglecting the sub-6-hourly wind variability and underestimated by about 70% without the sub-daily variability. Consistent with Zhai (2017) and D’Asaro (1985, 1995), the occurrence of winter storms induces larger inertial currents and *F_e_K_ea_* than during the summer.

**Figure 27:**
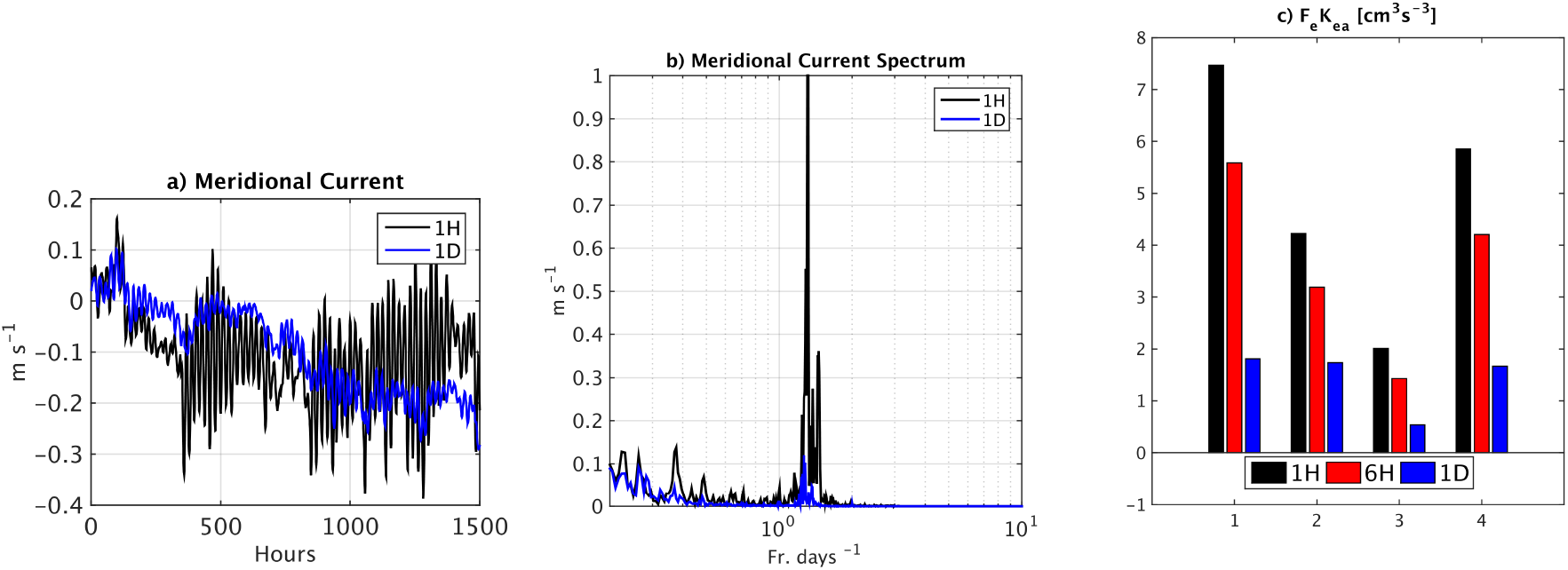
Surface current responses to hourly and daily wind forcing. (a) Hourly time series of the surface meridional current at (36°N, 122°W) (central California coast). The black and blue lines represent the simulation forced by the hourly-(1H, as in the USW4 simulation) and the daily-updated wind (1D). (b) The temporal spectrum of the surface meridional current from the 1H (black) and 1D (blue) simulations at (36°N, 122°W). (c) Mean ageostrophic eddy wind work (*F_e_ K_ea_*) (cm^3^ s^−3^) averaged between 30°N and 45°N and over a 500 km cross-shore distance from the coast in the 1H, 6H, and 1D simulations. As expected, the high-frequency wind forcing enhances the ageostrophic wind work and inertial currents.

## 6 Discussion

In this study regional atmospheric and oceanic model simulations are made for a 16-year hindcast period from 1995 to 2010. The simulations are evaluated against satellite and *in situ* measurements with an emphasis on the seasonal cycle and the mean and mesoscale circulations of the California Current System (CCS).

We evaluate the atmospheric forcing simulated by WRF and find, in general, a good agreement between the simulations and the measurements of the cloud cover, heat fluxes, and surface stress with modest discrepancies that are some combination of estimation errors and model biases. In particular, we show the ability of the atmospheric model to represent realistically the stratocumulus cloud deck in the northeastern Pacific. Then, by comparing the oceanic simulation to available measurements and previous modeling studies, we demonstrate the consistency of the simulations in representing the mean circulation and the seasonal and mesoscale variability of the CCS. Our results illustrate the benefits of using both oceanic and atmospheric regional simulations to simulate the seasonal variability of an eastern boundary upwelling system, at least in part because of the excessively coarse resolution in global models. Although some aspects of the interannual variability have been included in this study, more could be examined about low-frequency variability in the CCS.

The wind drop-off characteristics of a similar atmospheric simulation have been validated by Renault et al. (2016b). The simulation reported in this paper presents a good agreement with the measurements. These oceanic validations are also an indirect validation of the wind profiles simulated by WRF. An alternative simulation has been carried out using the CFSR reanalysis (Saha et al., 2010). Due to a poor representation of the wind drop-off, this simulation was characterized by an unrealistic poleward surface current and a poor representation of the mesoscale activity. The coarse resolution of CFSR (or other similar reanalysis) prevents using such a product to force this particular upwelling region and should not be used to investigate processes or trends, at least in the CCS.

Although not discussed here in any detail, the oceanic simulation is forced using various lateral open-ocean boundary forcing fields, such as Mercator or SODA. Differences in the lateral conditions can lead to significant changes in mean temperature and salinity (up to 0.5°C in SST and 0.5 PSU in *S*). Probably they are the primary cause for the the salinity biases present in the USW4 simulation, which are perhaps the largest inaccuracy of the simulation. We finally chose to force the simulations using Mercator with an additional mean monthly state correction toward the measurements from the World Ocean Database over the period 1995-2004. Nevertheless, uncer-tainty in open-ocean boundary conditions of the gyre-scale currents, density, and other water-mass properties do limit the possible accuracy of these quantities in a regional simulation.

An important contribution of this paper is our use over a long time period (1995-2010) of the parameterization of the wind and stress response to the current feedback suggested by Renault et al. (2016d) for the U. S. West Coast. Long-term comparisons with satellite measurements show realistic simulation results for the EKE and the energy transfer between the ocean and atmosphere with this feedback — and falsely large EKE values without it — in high-resolution models. Oceanic models, if uncoupled, should take into account the current feedback and by including a parameterization of the wind response such as Eq.(3) for a realistic kinetic energy transfer between the atmosphere and the ocean, and thus for a realistic level of mesoscale activity and mean circulation.

Finally, we discussed the importance of using a high-frequency wind forcing to represent the mean features of the CCS. In particular, consistent with Wu et al. (2016), we show the presence of high frequency wind prevents the use of monthly wind to force an oceanic model of the CCS. It leads to large errors in the mean stress and wind work inputs to the ocean, and, thus, to a poor representation of the mean and mesoscale currents. For the CCS we show that a 6-hourly wind forcing realistically represents the mean surface stress and the mean and mesoscale geostrophic currents. A daily wind forcing, such as QuikSCAT (commonly used to force an oceanic model), leads to an underestimation of the EKE by 7% for the CCS. However, a 1-hourly wind forcing is necessary for proper representation of the inertial currents.

In summary, we show the benefit of using both oceanic and atmospheric simulations for representing the mean physical state of the CCS. The atmospheric model is characterized by several biases such as too dry a lower atmosphere, too few clouds nearshore (although it realistically represents the stratocumulus cloud deck in the north of the domain), too much precipitation, and slightly too low a surface stress. As a response, the oceanic model has too large a surface salinity, too cold a SST, and too deep a MLD. Some other oceanic biases are controlled by the open-boundary conditions, such as the too-cold 150 m depth temperature. Perfect model-measurement agreement is an impossible goal both because of sampling limitations in model and observational data and because there are too many model design options and parameterization choices to ever be fundamentally correct or precise (McWilliams, 2007). Nevertheless, the USW4 system presented here is in fairly good overall agreement with the measurements that exist, and it has no glaring failures with respect to its primary behaviors. It thus provides a reliable physical foundation for assessing biogeochemical cycles and climate changes in the CCS (Deutsch et al., 2021a; Howard et al., 2020a, 2021; Kessouri et al., 2021b).

## Codes and Simulation Data

The physical and biogeochemical codes used for our simulations are at https://github.com/UCLA-ROMS/Code. The simulation model output archive data can be made available by an email request to the Corresponding Author.

## Acknowledgments

We appreciate support from the Office of Naval Research (ONR N00014-12-1-0939), the National Science Foundation (OCE-1419450), the California Ocean Protection Council grant (Integrated modeling assessments and projections for the California Current System), the Bureau of Ocean Energy Management, and the National Oceanic and Atmospheric Administration (DOC-NOAA NA15NOS4780186). Model simulations were carried out using the Extreme Science and Engineering Discovery Environment (XSEDE) and Yellowstone computers supported by the National Science Foundation at San Diego Supercomputer Center and NCAR, respectively. This study has been conducted using E.U. Copernicus Marine Service Information. MODIS level 2 data were downloaded from the NASA Web site (available at *http://ladsweb.nascom.nasa.gov*). GPCP data were provided by the NOAA/OAR/ESRL PSD, Boulder, Colorado, USA, from their web site at *https://www.esrl.noaa.gov/psd/*. Gliders data can be found at *https://spraydata.ucsd.edu/climCUGN/*

## Footnotes

2 In retrospect, a better result might have occurred had we done a density-space correction of the boundary mean *T* – *S* values, as is done for the biogeochemical properties (Deutsch et al., 2021a), but this is unlikely to overcome sampling error from data sparseness. A more elaborate procedure would be to adjust the boundary data to reduce interior bias, but this would be a form of data assimilation, which we otherwise have avoided.

## Notes

### Competing Interest Statement

The authors have declared no competing interest.

### Summary of Updates

This version of the manuscript has been revised following peer review.

